# Rewiring systemic nodulation control to engineer heat-tolerant nitrogen fixation in *Medicago truncatula* and chickpea

**DOI:** 10.64898/2026.10.01.755958

**Authors:** Manish Tiwari, Sairam Nagalla, Rakesh Kumar Swain, Swapnil Kumar, Valentina Infante, Junko Maeda, Manisha Yadav, Divya Mishra, Jean-Michel Ané

**Affiliations:** Department of Bacteriology, University of Wisconsin-Madison, Madison, WI 53706, USA; Department of Genetics, University of Delhi South Campus, New Delhi 110021, India; National Institute of Plant Genome Research, Aruna Asaf Ali Marg, New Delhi 110067, India; Department of Plant and Agroecosystem Sciences, University of Wisconsin-Madison, Madison, WI 53706, USA

## Abstract

Root nodule symbiosis allows legumes to meet their nitrogen requirements. However, the causes of symbiosis failure at high temperatures remain unknown. We demonstrated that elevated temperatures affect both the host and the microbial partner in *Medicago truncatula* and chickpea. Rhizobial viability and *nod* gene expression persisted under heat stress, but plant transcriptome analysis revealed a strong downregulation of the nodule organogenesis machinery. Nodule organogenesis was restored by constitutive overexpression of *NIN-LBD16-NFYA1-NFYB16*, *miR2111,* and *CLE5* knockdown but failed to restore nitrogen fixation. In contrast, the genetic manipulation of the competence pathway involving CEP peptides significantly improved both nodulation and nitrogen fixation. Constitutive loss of the autoregulation of nodulation (AON) pathway in the *SUNN* alleles also rescued root nodulation and nitrogen fixation. Therefore, heat stress perturbs the balance between AON and the competence pathway, and genetic manipulation of these pathways is a promising approach to engineer thermotolerant nitrogen fixation in crop legumes.

## Introduction

Symbiotic associations between plants and microbes such as rhizobia and mycorrhizal fungi help nutrient acquisition and provide abiotic stress tolerance^1^. Although nitrogen is one of the most abundant elements of the atmosphere (∼79%), its bioavailable forms, nitrate and ammonium, often limit plant growth and development. Therefore, modern agriculture largely depends on synthetic fertilizer application, which adds economic and environmental costs for farmers and the environment^2^. A sustainable alternative to reduce dependency on synthetic fertilizers is biological nitrogen fixation (BNF). Plants belonging to the nitrogen-fixing clade: Fabales, Fagales, Rosales, and Cucurbitales associate with diazotrophic bacteria in specialized root nodules to convert nitrogen into ammonia^2^. This association, known as root nodule symbiosis (RNS), enables legumes to meet their nitrogen requirements with minimal synthetic fertilizer input, but abiotic stresses such as drought, heat, and salinity impair this association. Investigations that resolve developmental checkpoint failures, or whether the impact is host- or microbe-derived, are lacking.

RNS can be broadly categorized into 5 phases: (i) early signaling involving lipochitooligosaccharides (LCOs) perception, (ii) infection involving pre-infection and infection thread formation, (iii) organogenesis, (iv) nitrogen-fixing function, and (v) regulation involving autoregulation of nodulation (AON). The host-rhizobia molecular dialogue begins when rhizobia recognize flavonoids released by the host plant, triggering the release of LCOs, also known as Nod factors. These LCOs are perceived by the plasma membrane receptors MtLYK3/LjNFR1 and MtNFP/LjNFR5 together with the coreceptor MtDMI2/LjSYMRK, activating a common symbiotic pathway that generates nuclear-localized calcium spiking, characterized by rhythmic oscillations in nuclear calcium concentration^3–6^. Nuclear calcium/calmodulin-dependent protein kinase MtDMI3/LjCCaMK decodes calcium signatures leading to expression of core transcriptional regulator NIN (NODULE INCEPTION)^7–10^. Downstream of LCO signaling, two independent processes are initiated: bacterial infection in the epidermal cell layer of the root, and nodule organogenesis by cell division of inner root cortical cells^11^, in both of which NIN acts as a master and essential regulator^12–14^. Although nodule organogenesis is normally associated with bacterial colonization, it can occur independently of bacterial infection through phytohormone action^15^. Cytokinin signaling mediated by CYTOKININ RESPONSE 1 (CRE1) kinase in inner-root cortical cells induces NIN expression^16^. NIN then co-opts the lateral root developmental program involving Lateral Boundary Domain16 (LBD16), NFYA1 and NFYB1 to drive nodule organogenesis, reflecting the extensive developmental overlap between the two processes^17,18^. Moreover, simultaneous overexpression of LBD16-NFYA1-NFYB1 is sufficient to rescue the nodule primordia-deficient phenotype of the *nin* mutant to that of infected nodules^17^.

Root nodule numbers are regulated by shoot-mediated systemic signaling involving receptors SUNN/NARK and CRA2, which bind CLE and CEP peptides respectively^19,20^. Consistent with SUNN/NARK’s role in autoregulation, mutant alleles *sunn-1, sunn-4*, and *lss* act in the shoot and result in a supernodulation phenotype with 4-10 times more nodules than wild-type in both *Medicago truncatula* and soybean (*Glycine max*)^21,22^. In contrast, hypernodulating mutants *rdn1* and *dar* affect genes acting in the root^23,24^. CLE peptides are synthesized in the root, transported to the shoot, and bind SUNN/NARK to activate AON^20^. In parallel, CEP peptides act through CRA2 to positively regulate nitrate acquisition, nitrogen fixation, mycorrhizal symbiosis and abiotic stress tolerance^19,25,26^. Thus, the AON and CEP-mediated competency pathways are conserved between Medicago and soybean, suggesting that Medicago mechanisms could translate to other legumes.

Rhizobia typically grow optimally at 25-30°C with effects on growth emerging above 35°C^27–29^. On the plant side, heat stress disrupts plant-microbe signaling and impairs root hairs with disintegrated infection threads, at 42°C^30^, delaying or abolishing infection and nodulation^31–33^. Heat stress during flowering and pod stages reduces soybean nodule biomass^34^, and causes peribacteroid membrane rupture^35^. Pea nodules show progressive senescence under heat stress^36,37^, and in alfalfa, up to 90% of nitrogen fixation was lost after 1 hour at 40°C or 4 hours at 36°C^38^. Thus, heat stress reduces nodule formation, nitrogen-fixing efficiency, and nodule longevity, yet the molecular basis spanning signaling, infection, organogenesis, and fixation remains unexplored, including whether impairment is host- or microbe-driven and whether signaling modules can be manipulated to restore symbiosis.

In this study, we characterized the effect of heat stress on RNS in Medicago and chickpea (*Cicer arietinum*) using developmental, genetic, physiological, and transcriptomic approaches, testing whether (i) heat stress affects both host and microbe, (ii) organogenesis can be rescued by the NIN-governed LBD16-NFYA1-NFYB16 module, and (iii) the AON pathway suppresses symbiosis under heat stress. We found that both host and microbial partners determine the heat stress response, with organogenesis and nitrogen fixation decoupled under heat stress, and identified AON as a key regulator whose manipulation can ameliorate heat stress effects on RNS.

## Results

### Heat stress impairs multiple stages of nodule symbiosis

To understand how heat stress affects RNS, we examined interactions between Medicago plants and *Sinorhizobium meliloti*, and between chickpea and *Mesorhizobium ciceri*. To determine the effect on early signaling, we monitored *ENOD11* expression using an *ENOD11*pro::*uidA* (GUS) reporter in wild-type Medicago plants inoculated with *S. meliloti*. Initially, GUS expression was confined to the rhizobial infection zone under both conditions, but expression subsequently progressed into mature nodules under control conditions, whereas under heat stress, expression remained localized to the roots **(Fig. 1a)**. Histochemical staining of stressed Medicago roots inoculated with *S. meliloti* carrying *hemA*pro::*lacZ* revealed infection threads arrested within the infection chambers or root hairs, with few transcellular infection threads progressing beyond the epidermis **(Fig. 1b-c)**. Consistent with blockage of infection thread development, roots formed few nodule primordia **(Fig. 1d-e)** and mature nodules at 5 and 14 days post inoculation (dpi), respectively **(Fig. 1f-g)**. In nodules of Medicago inoculated with a *S. meliloti* 1021 *nifH*pro*::uidA* reporter strain, *nifH* promoter activity was strongly evident throughout the mature nitrogen fixation zone under control conditions at 14 dpi, but was lost under stress **(Fig. 1h)**. The microscopic observation was consistent with the acetylene reduction assay (ARA) results, which showed nearly abolished nitrogen fixation at 14 dpi **(Fig. 1i)**. Together, these results indicate that heat stress disrupts RNS at multiple checkpoints, hampering the process from the early molecular dialogue between rhizobia and host through nitrogen fixation itself.

**Figure 1.**
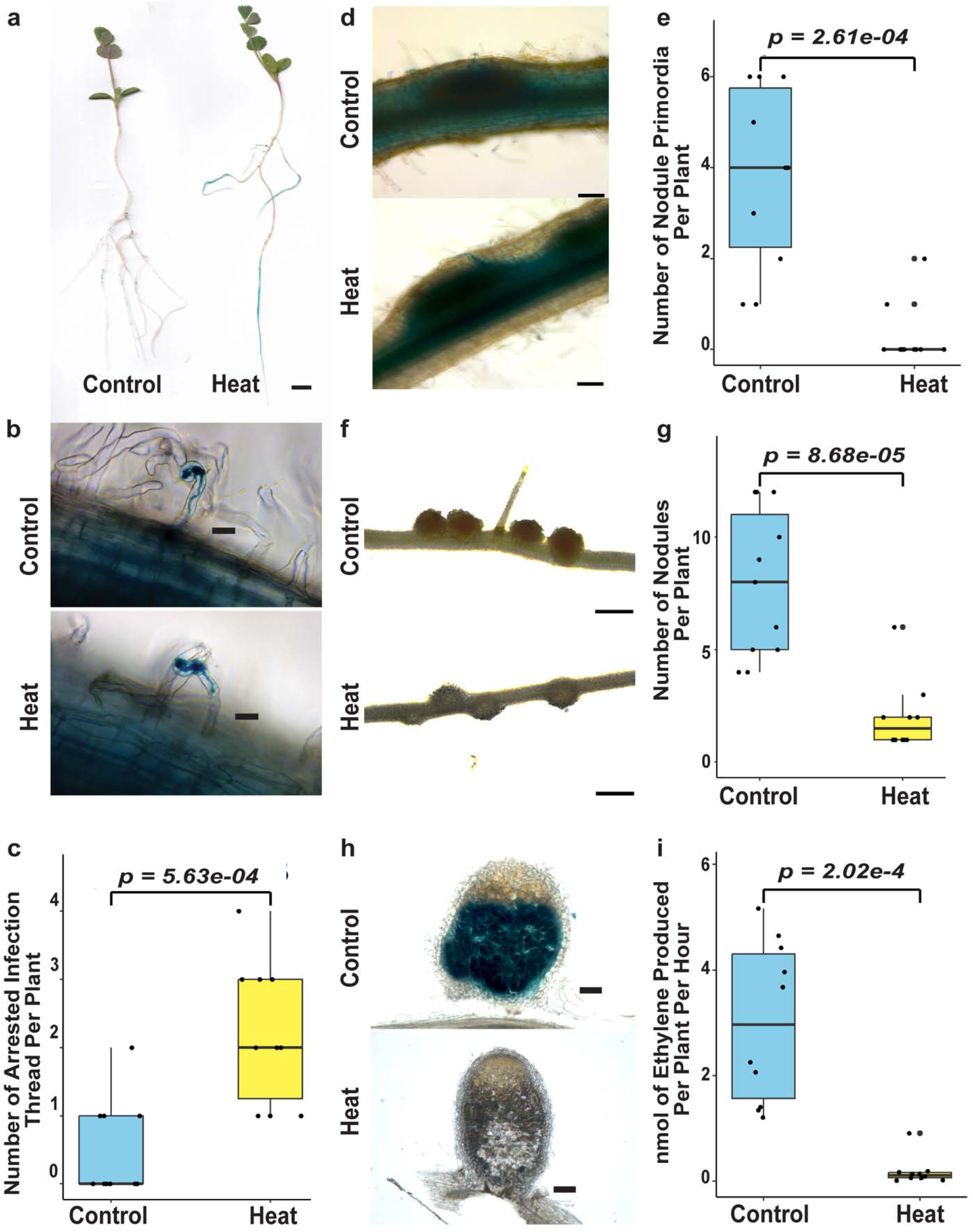
Heat stress impairs root nodule symbiosis in *M. truncatula*. **a,** Representative *ENOD11*pro::*uidA* expression in roots under control and heat stress conditions in wild-type plants containing *ENOD11*pro∷*uidA* (n = 10). **b,** Representative images at 3 dpi of infection threads formed by *S. meliloti* carrying *hemA*pro::*lacZ*, fully extended under control conditions and arrested under heat stress. **c,** Quantification of arrested infection threads at 3 dpi under control and heat stress conditions (n = 10). **d,** Representative nodule primordia at 5 days post inoculation (dpi). **e,** Quantification of nodule primordia at 5 dpi (n = 10). **f,** Representative nodules at 14 dpi under control and heat stress conditions. **g,** Quantification of nodule number at 14 dpi (n = 10). **h,** Functional nitrogen-fixing nodules at 14 dpi visualized by GUS staining following inoculation with *S. meliloti* 1021 *nifH*pro::*uidA* (n = 10 nodules)**. i,** Nitrogenase activity measured by the acetylene reduction assay (ARA) at 14 dpi (n = 10). Images correspond to 60-μm optical sections in (**h).** Scale bars: 1 cm (**a**), 50 μm (**b**) and 100 μm (**d,h**), 1 mm (**f**). Statistical significance was determined using two-sided Welch’s t-tests. Box plots show the median (center line), 25th and 75th percentiles (box), and whiskers extending to 1.5× the interquartile range, with individual data points shown as dots.

### Heat stress impairs rhizobial fitness and LCO output

Having established that heat stress affects multiple stages of RNS, we next sought to delineate its effects on the host versus the rhizobial partner. Growth curves of *S. meliloti* tested at three temperatures showed that the bacterium reached mid-exponential phase earliest (highest 1/T mid), produced the most biomass (highest carrying capacity, K), and had the largest area under the curve at 28°C. At the higher temperatures (30 and 32°C), the bacterium remained viable, but growth kinetics shifted. Compared to 28°C, exponential growth began later and the final biomass was lower, despite the maximum growth rate (μmax) being highest at 32°C. (**Extended Data Fig. 1a-e**). Similarly, a drop plate assay with *M. ciceri* also confirmed that the rhizobia survived well at higher temperatures with slower growth (**Extended Data Fig. 1f)**. Again, maximum *nodA* gene induction, measured using the *S. meliloti nodA*pro*::lacZ* reporter strain occurred at 28°C, which decreased as temperature increased to 32°C (**Extended Data Fig. 1g**). Interestingly, the *S. meliloti nodD*-overexpressing strain 6390 (LCO-overproducing, in the *S. meliloti* 2011 background) failed to rescue both nodule number and nitrogen fixation in Medicago **(Extended Data Fig. 1h–i)**. These results suggest that heat stress affects the rhizobial partner at multiple levels, including growth dynamics and *nod* gene expression. However, failure to rescue RNS with LCOs overproducing strain suggests that *nod* gene activation is not the primary reason for the defects in RNS.

### Transcriptomic landscape of heat-stressed nodulation reveals nuclear/signaling and stress-response signatures

To dissect the molecular mechanism behind defects in RNS, we performed RNA-seq on rhizobia inoculated Medicago roots under control and heat stress conditions across multiple time points **(Fig. 2)**. We observed extensive host transcriptome reprogramming, with the largest cohort of differentially expressed genes (DEGs) detected at 1 hour post-inoculation (hpi) **(Supplementary Table 1)**. Notably, Gene Ontology (GO) enrichment of the DEGs between control and heat stress tissues was dominated by abiotic stress, stimulus terms including ion, osmotic, salt and temperature and hormone, suggesting that heat stress response overshadows symbiotic signaling cascade and perturbs hormonal pathways **(Supplementary Fig. 1a)**. In the molecular function category, the genes were enriched for nucleotide-binding terms, ribosomal components and catalytic activity **(Supplementary Fig. 1b)**. Also, the cellular component category included terms related to the symplast, plasmodesmata, and cell-cell junctions **(Supplementary Fig. 1c)**. As the infection thread progression involves polarised growth directed by the nucleus and symplastic channeling between the epidermal and cortical cell layers, the enrichment of these terms correlates with the arrest of infection threads.

**Figure 2.**
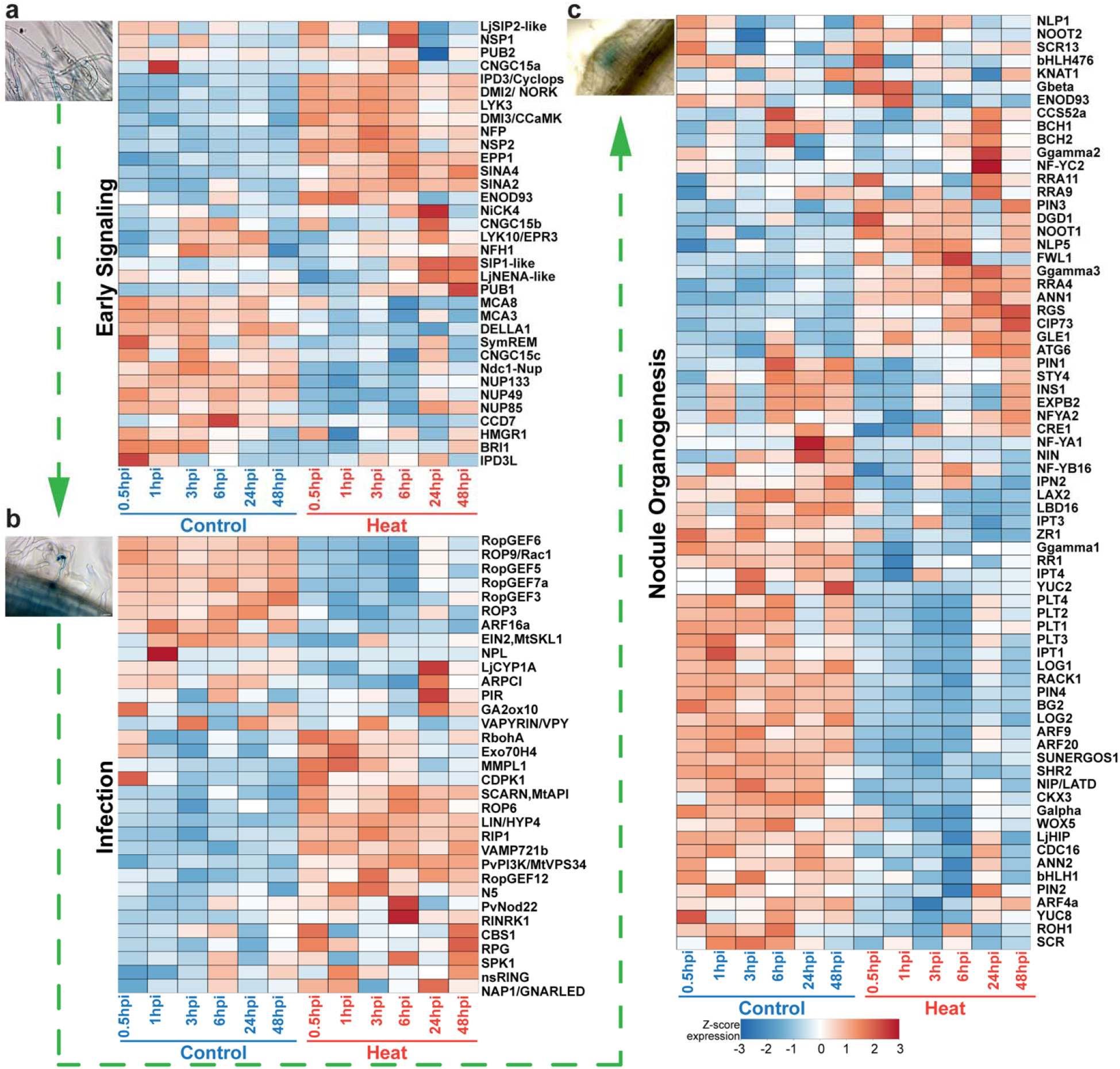
Transcriptomic analysis of the effects of heat stress on root nodule symbiosis in *M. truncatula*. **a,** Heatmaps showing the expression profiles of genes differentially expressed under control and heat stress conditions at 0.5, 1, 3, 6, 24, and 48 hours post-inoculation (hpi). Genes are grouped by their roles in early symbiotic signaling, **b,** infection, and **c,** nodule organogenesis. Insets show representative images of: **a,** early signaling, **b,** infection, and **c,** organogenesis. Data were obtained through RNAseq of *Medicago* roots inoculated with *S. meliloti* 1021, in three independent biological replicates under control and heat stress conditions at different time points. The color scale represents row-scaled Z-scores of log2 (TPM + 1) transformed gene expression clipped to a range of −3 to 3 shared across panels a-c.

We focused on how heat stress reshapes the expression trajectory of genes regulating early signaling, infection, and nodule organogenesis in Medicago **(Fig. 2a-c)**. Heat stress rewired the expression pattern across gene modules, with the infection and nodule organogenesis modules most strongly affected. Genes regulating rhizobial perception and calcium spiking activation, including the LCO receptors and co-receptor pair *LYK3* and *NFP, DMI2* and CSP activator *IPD3/CYCLOPS*, and downstream effectors were upregulated under heat stress. Contrastingly, the nuclear pore (NUP) complex subunits, *NUP133, NUP85, NUP49*, and *Ndc1-Nup* members, were downregulated, pointing to heat sensitivity of nuclear envelope machinery in Medicago **(Fig. 2a)**. These expression signatures indicate that rhizobial perception remains intact; however, the precise temporal window of calcium-signaling machinery activation is disrupted by loss of nuclear pore integrity. The defective timing of calcium-spiking activation disrupted downstream root nodule stages. The genetic module governing infection thread progression involving *ROP GTPases*, *VAPYRIN*, and *RopGEFs* were mostly downregulated, suggesting that cytoskeleton remodelling machinery is more heat-sensitive compared to the initial perception and root hair curling **(Fig. 2b)**. The organogenesis modules; cortical cell division and nodule primordium formation, including *NFYA1, NFYB16*, cytokinin biosynthesis/signaling genes *IPT1/3/4, LOG1/2,* auxin-related regulators *YUC2/8, ARF4a, PIN1,2,4, LAX2, PLT1/2/3/4*, and meristem identity genes *WOX5, SHR2, SCR*, were the most severely disrupted **(Fig. 2c)**. This data indicates that auxin-cytokinin cross-talk is a central node of heat sensitivity downstream of early signaling and infection.

### Restoring the *NIN-LBD16-NFYA1-NFYB16* organogenesis module rescues nodule number but not nitrogen fixation

Since heat stress suppressed *LBD16*, *NFYA1,* and *NFYB16,* an essential module downstream of *NIN* that also regulates lateral root organogenesis, we hypothesized that overexpressing a module comprising *NIN, LBD16*, *NFYA1,* and *NFYB16* together could potentially rescue nodule organogenesis. Medicago transgenic roots constitutively co-expressing this module did not show any increase in nodule number or nitrogen fixation under control conditions **(Extended Data Fig. 2a, c)**; however, a significantly increased nodule number relative to the vector control was observed under heat stress **(Extended Data Fig. 2b)**. Interestingly, the gain in nodule number under heat stress was not translated to symbiotic functions, as nitrogen fixation in the overexpressing lines and vector control was similar, indicating non-fixing nodule formation at 14 dpi **(Extended Data Fig. 2d)**. Therefore, the *NIN-LBD16-NFYA1-NFYB16* module successfully restored nodule organogenesis but failed at nitrogen fixation under heat stress. This result indicates that heat stress decouples nodule development from rhizobial accommodation inside the nodules, suggesting that a checkpoint operates beyond organogenesis, encompassing bacteroid differentiation and symbiosome maturation, to govern nitrogen fixation.

### CEP peptides are broadly downregulated under heat stress, and their exogenous/genetic restoration rescues nodulation

Our RNA-seq analysis showed heat-responsive repression of *CEP* genes **(Extended Data Fig. 3a)**. We hypothesized that supplementing CEP peptides could rescue heat-stress phenotypes. Based on the expression profile and difference in the C-terminal 15 amino acid region we synthesized 12 CEP peptides **(Fig. 3a)**. Exogenous application of eight of the twelve CEP peptides (all except CEP4, CEP9, CEP10 and CEP16) to Medicago roots significantly rescued nodule number compared to mock-treated roots. **(Fig. 3b)**. Based on the nodule numbers or the morphology of nodule structure we chose six different peptides (CEP1, 2, 5, 7, 8, 12) for further characterization in Medicago **(Extended Data Fig. 3b)**. Under heat stress, exogenous treatment of *ENOD11*pro::*uidA* wild-type with these peptides increased reporter expression in root nodules compared to mock treated plants **(Fig. 3c)**. The plants treated with all six selected CEP peptides in Medicago showed increased nodulation as well as a modest but significant recovery of nitrogen fixation for CEP1, 2, 5 and 7 **(Fig. 3d-e)**. Under heat stress, constitutive overexpression of these *CEP* genes in Medicago roots rescued nodule numbers for all six genes, but rescued nitrogen fixation only for *CEP1* and *CEP5* compared with the vector control **(Fig. 3f-g)**. CEP peptide treatment also improved overall plant growth, as shoot weight and root weight increased for all six peptides, and root length increased for CEP1, CEP5, and CEP7 **(Extended Data Fig. 3c-e)**.

**Figure 3.**
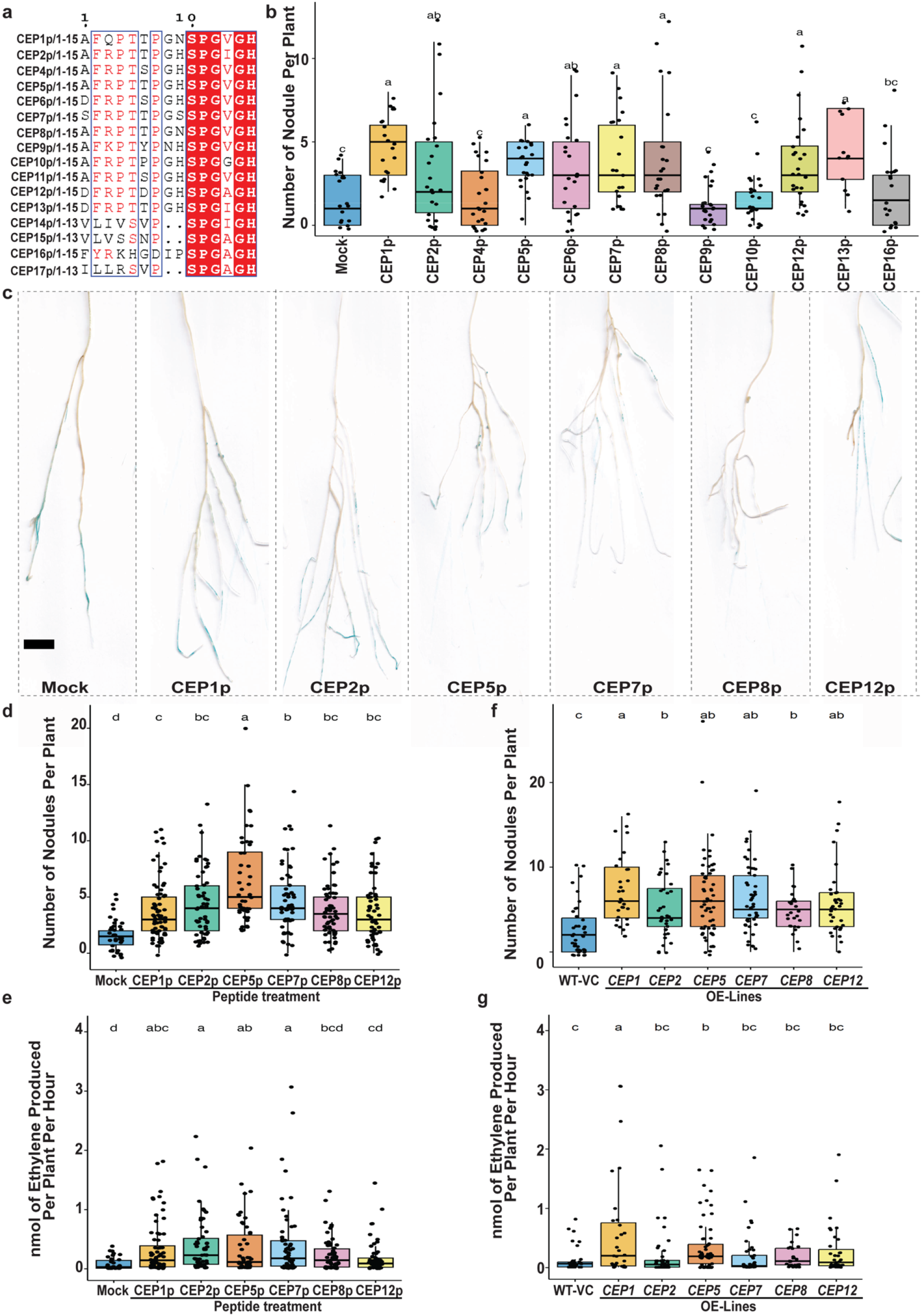
CEP peptides partially rescue nitrogen fixation under heat stress. **a,** Sequence alignment of the mature peptide regions of the 16 *M. truncatula* CEP family members. Red box indicates conserved amino acids across all of the CEP peptides. **b,** Quantification of nodule number in plants treated with 12 synthetic CEP peptides (1 μM) under heat stress compared with the mock-treated wild-type (WT) (Mock (n = 19), CEP1 (n = 21), CEP2 (n = 24), CEP4 (n = 24), CEP5 (n = 23), CEP6 (n = 23), CEP7 (n = 21), CEP8 (n = 24), CEP9 (n = 24), CEP10 (n = 23), CEP12 (n = 26), CEP13 (n = 12), CEP16 (n = 20)). **c,** Representative *ENOD11*pro::*uidA* expression in *M. truncatula* Jemalong A17 wild-type plants treated with CEP peptides or mock at 14 dpi under heat stress. **d,** Quantification of nodule number in heat-stressed plants treated with six selected CEP peptides (Mock (n = 36), CEP1 (n = 72), CEP2 (n = 56), CEP5 (n = 50), CEP7 (n = 60), CEP8 (n = 62), CEP12 (n = 56)). **e,** Nitrogenase activity measured by the ARA at 14 dpi (Mock (n = 36), CEP1 (n = 72), CEP2 (n = 56), CEP5 (n = 50), CEP7 (n = 60), CEP8 (n = 62), CEP12 (n = 56)). **f,** Quantification of nodule number in plants constitutively expressing selected *CEP* genes under heat stress compared with the empty-vector control (WT-VC (n = 41), *CEP1* (n = 31), *CEP2* (n = 35), *CEP5* (n = 61), *CEP7* (n = 49), *CEP8* (n = 27), *CEP12* (n = 35)). **g,** Nitrogenase activity measured by the ARA at 14 dpi in *CEP*-overexpressing plants. (WT-VC (n = 38), *CEP1* (n = 31), *CEP2* (n = 35), *CEP5* (n = 61), *CEP7* (n = 41), *CEP8* (n = 27), *CEP12* (n = 35)). Scale bars, 1 cm (**c**). Statistical significance was determined using one-way ANOVA followed by Fisher’s LSD test (p-value < 0.05). Different letters indicate statistically significant differences. Box plots show the median (center line), 25th and 75th percentiles (box), and whiskers extending to 1.5× the interquartile range, with individual data points shown as dots.

### The autoregulation of nodulation (AON) pathway is a target of heat stress, and its manipulation rescues root nodule number

After partially rescuing symbiosis via the CEP competence pathway, we turned to the AON pathway involving CLE peptides, the microRNA miR2111, and the LRR-RLK receptor kinase. Among *CLE* genes, *CLE5* showed higher expression during heat stress in Medicago and chickpea **(Supplementary Fig. 2, 6b)**. *CLE5* transcript expression increased under heat stress, and both plants treated exogenously with CLE5 and transgenic roots constitutively expressing *CLE5* showed greatly reduced nodule numbers (peptide treatment under control conditions; overexpression under control and heat stress conditions), confirming that CLE5 negatively regulates RNS **(Extended Data Fig. 4a-d)**. Plants with *CLE5*-RNAi showed a significant increase in nodule numbers but not in nitrogen fixation, as measured by ARA **(Fig. 4a-c)**. Similarly, overexpression of the systemic AON antagonist *miR2111* increased nodule numbers under both control and heat stress conditions; however, unfortunately, no recovery of nitrogen fixation was observed **(Fig. 4d-f, Extended Data Fig. 4e)**. Therefore, transiently relieving AON rescues nodule number but fails at nitrogen fixation, pointing again towards a decoupling of organogenesis from nitrogen fixation under heat stress.

**Figure 4.**
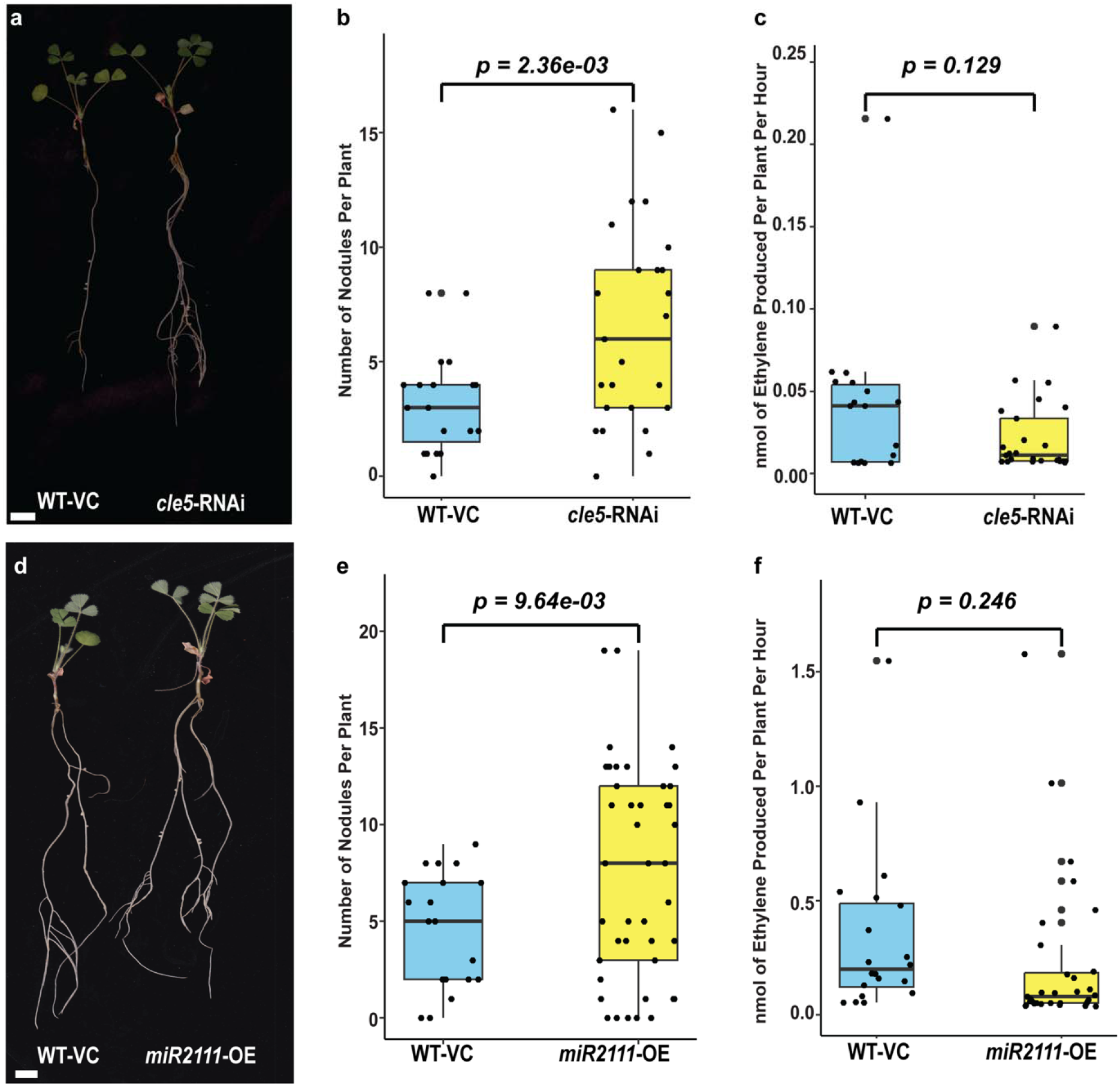
The autoregulation of nodulation (AON) pathway regulates root nodule symbiosis under heat stress. **a,** Representative images of *CLE5* knockdown plants compared to vector control under heat stress conditions. **b,** Quantification of the number of nodules in *CLE5* knockdown plants compared to vector control under heat stress conditions (WT-VC (n = 19), *CLE5*-RNAi (n = 25)). **c,** Nitrogenase activity measured by the ARA at 14 dpi under heat stress conditions (WT-VC (n = 18), *CLE5*-RNAi (n = 25)). **d,** Representative images of miR2111 overexpression plants compared to vector control under heat stress conditions. **e,** Quantification of the number of nodules in *miR2111* overexpression plants compared to vector control under heat stress conditions (WT-VC (n = 20), *miR2111-*OE (n = 42)). **f,** Nitrogenase activity measured by the ARA at 14 dpi under heat stress conditions (WT-VC (n = 20), *miR2111-*OE (n = 31)). Scale bars, 1 cm (**a, d**). Statistical significance was determined using two-sided Welch’s t-tests. Box plots show the median (center line), 25th and 75th percentiles (box) and whiskers extending to 1.5× the interquartile range, with individual data points shown as dots.

### Constitutive hypernodulation mutants show superior tolerance to heat stress at the whole-plant level

Because the competence and AON pathways helped plants tolerate heat stress, we hypothesized that complete loss of the AON pathway could rescue both nodule numbers and nitrogen fixation. To investigate how AON mutants respond to heat stress, we used hypernodulation mutants: *sunn-1, sunn-4, lss, dar1-1,* and *rdn1-2*. Under heat stress conditions, hypernodulation mutants showed a better overall phenotype in terms of plant biomass than wild-type plants **(Fig. 5a-b)**. *lss* and *sunn-4* formed ∼18 fold more nodules than wild-type plants under heat stress, while *sunn-1, dar1-1* and *rdn1-2* formed ∼5-8 fold more **(Fig. 5c)**. The increase in nodule numbers translated into functional recovery as *lss* mutants fixed nitrogen under heat stress comparable to the wild-type plants under control conditions **(Fig. 5d)**. Heat stress did not reduce shoot weight in wild-type plants, but *lss* gained shoot weight under heat stress **(Supplementary Fig. 3a)**. Root weight increased under heat stress in *sunn-1* and *dar1-1* **(Supplementary Fig. 3b)**. Root length increased under heat stress in wild-type, *sunn-1*, *sunn-4*, and *lss* plants but decreased in *rdn1-2* **(Supplementary Fig. 3c)**.

**Figure 5.**
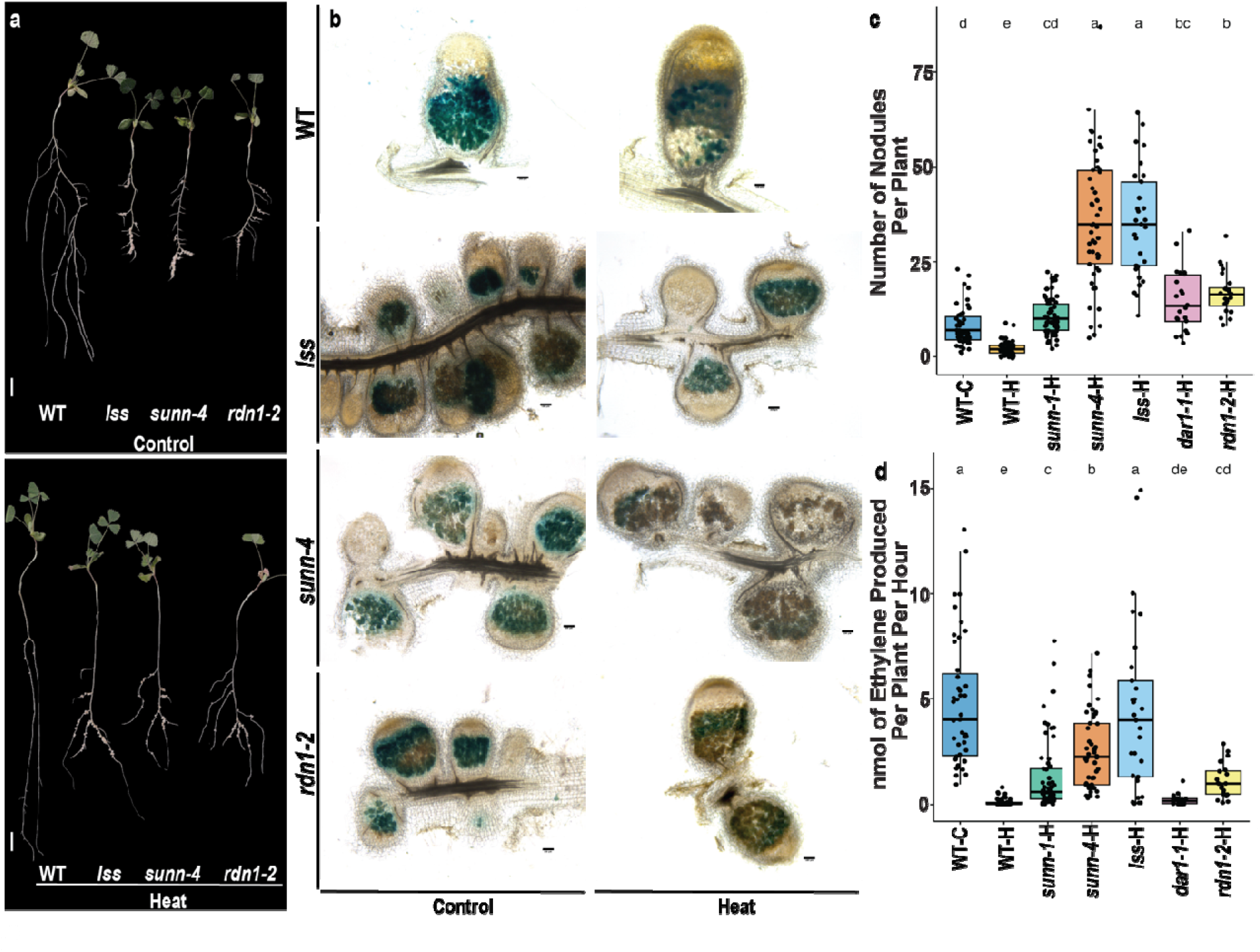
*M. truncatula* hypernodulation mutants tolerate heat stress better than wild-type plants. **a,** Representative images of *M. truncatula* hypernodulation mutants compared to wild-type (WT) under control and heat stress conditions. Two additional hypernodulation mutants, *sunn-1* and *dar-1,* were included only in the quantitative assays shown in **c** and **d. b,** Nitrogen-fixing nodules of wild-type, *lss, sunn-4, rdn1-2* under control and heat stress conditions visualized by GUS staining following inoculation with *S. meliloti* 1021 *nifH*pro::*uidA* (n = 10 nodules). **c,** Quantification of the number of nodules in wild-type under control and heat stress, and *sunn-1, sunn-4, lss, dar-1,* and *rdn1-2* under heat stress conditions. (WT-C (n = 43), WT-H (n = 54), *sunn-1* (n = 61), *sunn-4* (n = 50), *lss* (n = 29), *dar-1* (n = 22), *rdn1-2* (n = 24)). **d,** Nitrogenase activity measured by the ARA at 14 dpi under heat stress conditions. (WT-C (n = 43), WT-H (n = 53), *sunn-1* (n = 61), *sunn-4* (n = 50), *lss* (n = 29), *dar-1* (n = 22), *rdn1-2* (n = 24)). Scale bars: 1 cm (**a**) and 100 μm (**b**). Images correspond to 60-μm optical sections in (**b**). Statistical significance was determined using one-way ANOVA followed by Fisher’s LSD test (p-value < 0.05). Different letters indicate statistically significant differences. Box plots show the median (center line), 25th and 75th percentiles (box), and whiskers extending to 1.5× the interquartile range, with individual data points shown as dots.

### The effect of heat stress and rescue strategies translates to chickpea

To test conservation, across an agronomically important legume, chickpea, we investigated the effect of heat stress on chickpea RNS. Chickpea plants also showed a significant reduction in root nodule numbers **(Fig. 6a-b)**. Further confocal imaging of chickpea nodules stained with SYTO-13 showed dense and uniform fluorescence under control conditions, reflecting tightly packed infected cells, versus reduced and patchy fluorescence, representing disorganized rhizobial distribution under heat stress **(Fig. 6c)**. Chickpea RNA-seq analysis showed a distinct expression pattern dominated by a time-dependent manner instead of a clear demarcation based on conditions observed in Medicago **(Supplementary Fig. 4a-c, Supplementary Table 2)**. The differences in symbiotic gene expression profiles between Medicago and chickpea suggest species-specific sensitivity of RNS to heat stress. Overall, GO terms related to stress-response signatures, ribosome/translation, catalytic activity and plasmodesmata, and cell-cell junctions were shared between Medicago and chickpea **(Supplementary Fig. 5a-c)**.

**Figure 6.**
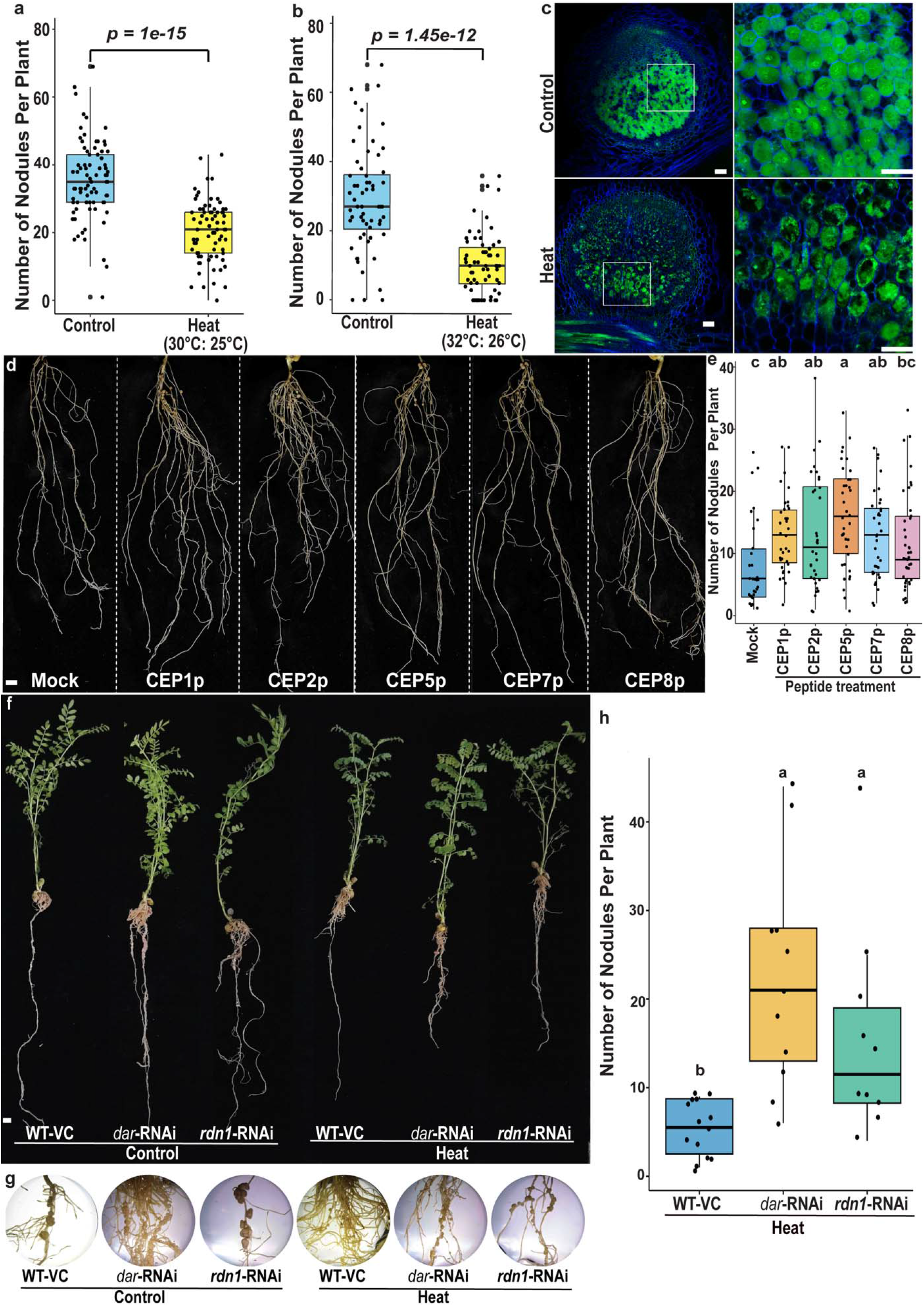
The effect of heat stress on root nodule symbiosis is conserved in *Cicer arietinum* (chickpea). **a,** Quantification of nodule number in chickpea at 30°C (Control (n = 78), Heat (n = 75)) and **b,** 32°C (Control (n = 60), Heat (n = 60)). **c,** Confocal images showing rhizobia stained with SYTO13 during chickpea root nodule development (n = 14 nodules). **d,** Representative images of heat-stressed chickpea plants treated with five selected CEP peptides or mock. **e,** Quantification of nodule number in heat-stressed chickpea plants treated with five selected CEP peptides compared with the mock-treated control. (Mock (n = 30), CEP1 (n = 35), CEP2 (n = 34), CEP5 (n = 37), CEP7 (n = 36), CEP8 (n = 40)). **f,** Representative images of chickpea empty vector plants, *dar*-RNAi and *rdn1-RNAi* lines under control and heat stress conditions. **g,** Inset images of nodules formed on empty vector, *dar*-RNAi and *rdn1*-RNAi lines under control and heat stress conditions. **h,** Quantification of the number of nodules in empty vector (n = 14), *dar*-RNAi (n = 11) and *rdn1*-RNAi (n = 10) lines under heat stress conditions. Images correspond to 60-μm optical sections in (**c**). Scale bars, 100 μm (**c**) and 1 cm (**d,f**). Statistical significance was determined using two-sided Welch’s t-tests in **a** and **b**. Statistical significance was determined using one-way ANOVA followed by Fisher’s LSD test (p-value < 0.05) in **e** and **h**. Different letters indicate statistically significant differences. Box plots show the median (center line), 25th and 75th percentiles (box), and whiskers extending to 1.5× the interquartile range, with individual data points shown as dots.

Altogether, the transcriptional profiling revealed that heat stress disrupts early signaling synchronization, disrupts infection thread progression, and causes an almost complete collapse of late organogenesis, as observed across legumes. Further, we identified 14 *CEP* genes in the chickpea genome **(Supplementary Table 3).** Out of these, six *CEPs* were orthologous to Medicago *CEPs* characterized in this study (*CEP1, CEP2, CEP5, CEP7, CEP8,* and *CEP12*). RNA-seq analysis showed heat-responsive repression of the *CEP* genes (*CEP1, CEP2, CEP4, CEP5, CEP7, CEP8* and *CEP12*) in chickpea, indicating that CEP suppression is conserved between legume species **(Supplementary Fig. 6a)**. We treated chickpea with five of these peptides (CEP1p, CEP2p, CEP5p, CEP7p, and CEP8p) to verify whether CEPs can ameliorate the effect of heat stress across legumes. Although the chickpea CEP12 protein showed similarity to Medicago CEP12, it lacked the conserved functional 15-amino-acid regions. CEP1p, CEP2p, CEP5p, and CEP7p significantly improved chickpea nodule numbers under heat stress compared with the mock treatment, whereas CEP8p did not **(Fig. 6d-e)**. Together, these results show that CEP peptides promote symbiotic competence, resulting in partial rescue of RNS across legumes. To test whether the heat-stress rescue phenotype is conserved in chickpea, we generated *dar* and *rdn1 RNAi lines.* Both RNAi lines showed significantly increased nodule number under heat stress compared with the vector control **(Fig. 6f-h).**

## Discussion

This study shows that heat stress affects RNS at several stages, including early signaling, infection, organogenesis, and nitrogen fixation. The defects are both host- and rhizobial partner-dependent, as rhizobial growth and *nod* gene activity are affected. Interestingly, enhancing plant-microbe signaling via LCOs overproduction could not rescue the phenotypes, arguing against the hypothesis that enough bacterial signals were absent **(Extended Data Fig. 1)**. In fact, the early signaling event was enhanced, as evidenced by transcriptomic signatures of early perception suggesting proper LCOs perception. Therefore, reduced bacterial fitness could play a major role in the observed heat stress phenotypes, and further investigations are needed to fully decipher microbial aspects of the stress response. The earliest disturbance of the common symbiotic pathway occurs in nuclear pore complex members and downstream signaling, which are downregulated **(Fig. 2)**. In our previous study, we showed that nucleoporins regulate localization of the calcium channels Castor and Pollux in *Lotus japonicus* and DMI1 in Medicago on the inner nuclear membrane, and their reduced expression changes localization of ion channels from the inner nuclear membrane to the outer nuclear membrane^39^. Therefore, low nucleoporin expression might disrupt nuclear-localized calcium spiking under heat stress, identifying this as an early heat-responsive checkpoint for symbiosis. The dominance of stress-related GO terms over symbiosis under heat stress in both Medicago and chickpea suggests a possible competition of resource allocation, in which hosts prioritize stress tolerance over symbiotic developmental programs^40^. The RNA-seq analysis in Medicago and chickpea also highlighted similarities and differences in expression patterns, which could reflect both host and microbial survival and growth^41^. A dual RNA-seq of host and symbiont is needed to discern host- and microbe-driven regulations of symbiosis under stress.

Heat stress perturbs the genetic circuitry regulating nodule organogenesis, primarily through auxin-cytokinin crosstalk. NIN orchestrates lateral root organogenic circuitry, acting through LBD16 and the NFYs to promote nodule primordium development^17,18^. Because heat stress reduced expression of these genes, we reasoned that restoring them could rescue nodule organogenesis. Overexpression of the *NIN-LBD16-NFYA1-NFYB16* module rescued nodule number but not nitrogen fixation **(Extended Data Fig. 2)**, indicating that fully functional nodule development requires a synchronous coupling of molecular, cellular, and biochemical events that heat stress evidently disrupts, decoupling organogenesis from nitrogen fixation and preventing mature symbiosis. This decoupling is consistent with the distinct developmental roles of the module’s components. NIN, a cytokinin-inducible master regulator of symbiosis, acts early after rhizobial perception to induce LBD16, which promotes localized auxin biosynthesis via STYLISH and YUCCA, engaging lateral-root organogenic circuitry to initiate nodule primordia. NIN also activates the NFYs, which govern rhizobial infection and cortical cell division during nodule structure formation^17,42^. Together, the module drives organogenesis and infection thread progression into dividing cortical cells, without itself contributing to nitrogen fixation.

A similar pattern emerged from *CLE5* RNAi and *miR2111* overexpression, important players in the AON pathway^19,20^. Both increased nodule number by interfering with the shoot-derived AON checkpoint, yet converged on the same outcome as the organogenesis module: more nodules without better nitrogen fixation **(Fig. 4)**. Those 3 independent genetic interventions *NIN-LBD16-NFYA1-NFYB16* overexpression, *CLE5* RNAi, and *miR2111* overexpression each rescued nodule number while failing to rescue nitrogen fixation indicates that nodule organogenesis and nitrogen fixation are governed by genetically separable programs under heat stress.

CEP peptides, a part of a competence pathway, are synthesized by the roots under limited nitrogen conditions and regulate systemic nitrogen demand signaling^43,44^. Multiple CEPs showed dynamic expression changes under heat stress in Medicago and chickpea, suggesting an overlap between CEP signaling and the heat stress response. CEP treatment and overexpression enhanced nodulation with a modest gain in nitrogen fixation and overall plant health, where an increase in shoot weight, root weight, and root length was observed **(Extended Data Fig. 3c-e)**. Overall, CEPs treatment suggests a general growth-promoting effect that benefits the symbiosis machinery, consistent with their demonstrated role in integrating nitrogen status with root architecture. The CEPs regulatory module remains conserved across legumes, as orthologous CaCEP peptide treatment increased nodule numbers under stress **(Fig. 6d-e)**, complementing heat-responsive CaCEP expression dynamics. These results lay the foundation for the translational relevance of CEP peptide-mediated heat stress tolerance response to the food legume crop chickpea beyond the model plant Medicago.

The genetic manipulations involving NIN-LBD16-NFYA1-NFYB16, CLE5/miR2111, and CEP peptides achieved only partial rescue under heat stress; therefore, we tested hypernodulation mutants producing more than ∼10-fold nodulation in plants. AON genes inhibit nodulation systemically and pleiotropically through long-distance root-shoot signaling, and disrupting this AON pathway conferred tolerance to symbiosis under heat stress. Heat stress profoundly inhibited nodulation and nitrogen fixation in wild-type plants; however, *lss* mutants fixed nitrogen at the level of wild-type plants under control conditions, and *sunn-4* mutants reached the next highest level. ARA activity and *nifHpro::uidA* staining confirmed that the hypernodulation mutants had a persistent nitrogen fixation zone, a result attributed to increased nodule numbers and to the formation of functional nodules. The protection mechanism was also conserved in chickpea, where *dar*-RNAi and *rdn1*-RNAi lines showed significantly higher nodulation compared to vector controls under heat **(Fig. 6f-h)**, indicating conservation across species. Recovery of the heat-stressed symbiotic phenotype was best in *lss* and *sunn-4* compared to *sunn-1, rdn1-2,* and *dar1-1*, suggesting that the extent of AON disruption dictates thermotolerance, consistent with *sunn-1*, *sunn-4*, and *lss* representing increasingly severe losses of *SUNN* function. Moreover, the *SUNN* alleles (*lss* and *sunn-4*), which act in the shoot, showed better tolerance than the root-acting mutants (*rdn1-2* and *dar1-1*), suggesting that shoot-derived signals also control the heat stress response of the root. However, this comparison rests on a single shoot-acting gene. Because AON pathway genes respond to nitrogen-carbon balance, their disruption may have shifted the balance toward symbiosis under stress^20,45^.

Several questions remain unaddressed, including the role of rhizobia and the mechanism underlying symbiosome deformation under heat stress, involving leghemoglobin function, oxygen protection, and NCR peptide-mediated bacteroid differentiation. Strategies to move beyond organogenesis toward improved nitrogen fixation, potentially through plant carbon partitioning under stress, also warrant follow-up. In conclusion, this work identifies the systemic autoregulatory pathway as the most promising target for engineering thermotolerant symbiosis.

## Methods

### Plant material and growth conditions

*Medicago truncatula* wild type (WT) Jemalong A17 and the *sunn-1, sunn-4, lss, rdn1-2,* and *dar1-1* mutants were scarified in H₂SO₄, surface sterilized with bleach, stratified on 1% agar plates at 4°C for two days, and germinated at room temperature for one day. Germinated seedlings were transferred to CYG growth pouches supplemented with modified Fahraeus medium and grown at 23/18°C (16 h light/8 h dark). 3-day-old seedlings were inoculated with *Sinorhizobium meliloti* at OD_600_ = 0.01 and transferred either to control conditions (23/18°C) or heat stress (30/25°C). We supplied modified Fahraeus medium every three days to prevent desiccation stress. Chickpea *(Cicer arietinum)* cultivar BGD256 seeds were surface sterilized with 2% sodium hypochlorite for 3 min, washed 8-10 times with sterile water, and germinated on autoclaved germination paper. We transferred two-day-old seedlings to germination pouches containing nitrogen-free McKnight’s solution for overnight acclimation at 23°C (16 h light/8 h dark). Three-day-old seedlings were inoculated by dipping the roots in 0.1 OD *Mesorhizobium ciceri* CC1192 for 45 min and immediately transferred to nitrogen-free soil supplemented with nitrogen-free McKnight’s solution. Plants were grown under 23/18°C (control), 30/25°C, or 32/26°C (heat stress) (16 h light/8 h dark).

### Bacterial culture and growth assays

*S. meliloti* 1021, *S. meliloti* 2011 and its *nodD*-overexpressing derivative 6390, and *M. ciceri* CC1192 were maintained on freshly streaked tryptone yeast (TY) agar plates. Primary cultures were grown for two days, followed by overnight secondary cultures. Cultures were adjusted to the required optical density before plant inoculation. For liquid growth assays, cultures were adjusted to OD_600_ = 0.7, diluted to a final OD_600_ = 0.01 in TY broth medium in 96-well plates, covered with breathable film, and incubated at 28, 30, and 32°C in an Agilent BioTek Epoch 2 plate reader for 96 h. Each temperature was run once on one 96-well plate with 72 technical replicate wells and 24 blank wells. For plate growth assays, secondary cultures were grown until OD_600_ = 1, serially diluted to OD_600_ = 0.1 and 0.05, and 5 μl of each dilution was spotted onto yeast mannitol broth (YMB) agar plates. Plates were incubated at 28, 30, and 32°C. *S. meliloti* GMI5874 carrying a *nodA*pro-*lacZ* transcriptional fusion^46^ was grown overnight in Vincent minimal medium (VMM) at 28°C. Cultures were diluted into fresh VMM to an initial OD₆₀₀ of 0.004 and supplemented with 10 μM luteolin or an equivalent volume of DMSO as a mock treatment. Cultures were incubated at 28, 30, or 32°C with shaking at 200 r.p.m. and harvested at an OD₆₀₀ of 0.2-0.7. β-galactosidase activity was measured using the ONPG-based liquid assay adapted from^47^. Briefly, cells from 2 mL of culture were collected, resuspended in Z buffer, permeabilized with chloroform, and assayed using 4 mg mL⁻¹ ONPG. Reactions were performed at 30°C and terminated with Na₂CO₃. The OD₄₂₀ and OD₅₅₀ were measured, and β-galactosidase activity was expressed as Miller units. Eight biological replicates were analyzed for each temperature × treatment combination.

### Identification of *CEP* and *CLE* genes in chickpea

The chickpea genome lacks annotated CEP and CLE genes; therefore, *M. truncatula* CEP protein and CDS sequences were used as queries in TBLASTN and BLASTP searches against the chickpea genome (*C. arietinum* ICC 4958, desi type, whole genome v3 https://www.pulsedb.org/organism/641). Only 14 CEPs were identified in chickpea. For CLE gene identification, we performed TBLASTN and BLASTP searches and additionally curated the CLE genes previously identified by Gentile *et al.* (2025)^48^.

### RNA sequencing and analysis

Roots of Medicago and chickpea inoculated with *S. meliloti* and *M. ciceri*, respectively, were harvested under control and heat stress conditions at 0.5, 1, 3, 6, 24, and 48 hours post inoculation (hpi). Total RNA was extracted using the Qiagen RNeasy Plant Mini Kit followed by DNase treatment. Paired-end RNA sequencing was performed on the Illumina platform from three biological replicates for each time point. The raw FASTQ reads were adapter-trimmed and filtered for Q30 reads with the FastP program. SortMeRNA was used to remove non-coding RNA reads before proceeding with transcript quantification using Kallisto with 100 bootstrap values. Sleuth program in R was used to determine differentially expressed genes using the Wald test and statistical significance at p-value ≤ 0.05 and false-discovery rate (q-value) ≤ 0.01^49^. GO enrichment analysis was performed using the Plant Transcriptional Regulatory Map database with a threshold p-value ≤ 0.01^50^.

### Constructs generation

Golden Gate cloning was used to generate overexpression constructs for *NIN-LBD16-NFYA1-NFYB16* driven by the *Lotus japonicus* ubiquitin promoter, *miR2111*, *CEP* genes, and *CLE5* driven by the CaMV35S promoter, as described in^51^. Individual domesticated Level 0 DNA sequences for Golden Gate cloning were synthesized by Synbio Technologies.

For RNAi constructs, the CDS of Medicago *CLE5* and chickpea *DAR* and *RDN1* were retrieved from the NCBI database. siRNA candidates were selected using pssRNAit. RNA was extracted from 3-day-old Medicago and chickpea roots inoculated with *S. meliloti* and *M. ciceri*, respectively, using the Direct-zol RNA Microprep Kit according to the manufacturer’s protocol. cDNA was synthesized from 1 μg of RNA using the Verso cDNA Synthesis Kit. The selected siRNA candidates were amplified using the primers listed in **Supplementary Table 4**. Entry clones were generated using the pENTR/D-TOPO cloning kit (Invitrogen) and recombined into the pK7WG2 destination vector using LR Clonase (Invitrogen).

### *Agrobacterium rhizogenes*-based root transformation

For *Medicago*, *Agrobacterium rhizogenes MSU440* carrying overexpression constructs for *CEP* genes, *miR2111, NIN-LBD16-NFYA1-NFYB16,* and *CLE5* overexpression and RNAi was used for root transformation as described in^39^. All constructs contained a TdTomato transformation marker driven by the Arabidopsis ubiquitin promoter to screen transformed roots. For chickpea, *A. rhizogenes* strain R1000 carrying the *dar* and *rdn1* RNAi constructs was grown on TYA plates with appropriate antibiotics. Germinated Medicago wild-type or chickpea BGD256 seedlings were wounded at the hypocotyl, inoculated with *Agrobacterium*, and transferred to Fahraeus medium. Transgenic roots were identified by Tdtomato/DsRed fluorescence; non-transformed roots were removed, and composite plants were used for nodulation experiments.

### Peptide treatments

The conserved 15-amino acid C-terminal regions of CEP peptides were synthesized (Biomatik Corporation, ≥95% purity). Three-day-old Medicago seedlings were treated with 10⁻ M peptide by applying 5 mL peptide solution to each growth pouch for 45 min before inoculation with 0.01 OD_600_ *S. meliloti* and transfer to control or heat stress conditions. For chickpea, synthetic CaCEP1, CEP2, CEP5, CEP7, and CEP8 peptides were synthesized (Jetta Meditech, ≥95% purity), and two-day-old seedlings were treated overnight with 1 μM peptide before inoculation with 0.1 OD_600_ *M. ciceri* CC1192 and transferred to nitrogen-free sand supplied with nitrogen-free McKnight’s solution. The phenotype for nodule numbers and nitrogen fixation was assessed at 14 dpi.

### Acetylene reduction assays

At 14 dpi, roots were harvested into 5 mL headspace GC vials, and 1 mL of air was replaced with 1 mL of acetylene gas. Vials were incubated overnight in the dark at room temperature, and ethylene production was measured using a Shimadzu gas chromatograph. Values were normalized to nmol per plant per hour.

### Microscopic analysis of infection threads and nodules

Three-day-old *Medicago* seedlings were inoculated with *hemA*pro::*lacZ*-expressing *S. meliloti*. Plants were maintained under control or heat stress conditions for 3 and 5 days, and infection threads and nodule primordia were quantified as described in^52,53^. Nitrogen-fixing bacteroids were visualized using *S. meliloti* carrying a *nifH*pro*::uidA* fusion in 60 μm nodule sections as described in^53,54^. For live bacterial cell imaging, thin chickpea nodule sections were stained with SYTO13 and calcofluor white, followed by confocal microscopy to determine the live-to-dead cell ratio under control and heat stress conditions^53^.

### Statistical analyses

Two-sided Welch’s *t*-test was used for comparison between two genotypes. When analyzing multiple genotypes or conditions, one-way analysis of variance (ANOVA), followed by Fisher’s least significant difference (LSD) test for pairwise comparisons, was used. Statistical significance was assessed at p-value < 0.05. Data are presented as individual biological observations, with box plots showing the distribution of the measurements and individual data points overlaid. For the rhizobial growth curve assay, a Gompertz growth model was fit to each well’s OD_600_ trajectory by nonlinear least squares (nlsLM, minpack.lm v1.2.4) in R v4.4.1 to extract the maximum growth rate (μ_max_, the steepest slope of the fitted curve), time to mid-exponential phase (T_mid_, reported as 1/Tmid so that higher values indicate faster growth), carrying capacity (K, the asymptote of the fitted curve), and area under the fitted curve over 96 h (AUC)^55^. Groups were compared using a Kruskal-Wallis test followed by Dunn’s post-hoc test (Benjamini-Hochberg-corrected) for μ_max_, 1/T_mid_, and K, and a Welch ANOVA followed by the Games-Howell post-hoc test for AUC, using rstatix v0.7.2. Letters in the figures show compact letter displays from the post-hoc comparisons (α = 0.05). For the *nodA* gene induction assay, Miller-unit values were log-transformed before analysis. The six temperature × treatment combinations were analyzed as a single factor using one-way ANOVA in R version 4.5.3. Residual normality was assessed using the Shapiro-Wilk test, and homogeneity of variance was assessed using Levene’s test. Because residuals were normally distributed but variances were unequal among groups, pairwise comparisons were performed using Dunnett’s T3 test, which does not assume homogeneity of variance. Statistical significance was defined as p-value < 0.05.

### Reporting summary

Further information on research design is available in the Nature Portfolio Reporting Summary linked to this article.

## Data availability

RNA-seq data sets are available via the National Center for Biotechnology Information Sequence Read Archive under accession number PRJNA1520955 at http://www.ncbi.nlm.nih.gov/sra/. Source data are provided with this paper.

## Acknowledgements

We thank Prof. Julia Frugoli for providing the seeds of *sunn-4, lss, rdn1-2,* and *dar1-1*

Medicago mutants.

## Author contributions

M.T., D.M., and J.M.A. designed and supervised the study and revised the manuscript. M.T., R.K.S., S.N., S.K., J.M., and V.I. performed the experiments. R.K.S., S.K., and D.M. performed all chickpea and *Mesorhizobium ciceri-related* experiments. S.N. and R.K.S. helped M.T. in acquiring Medicago phenotypic data. V.I. and M.T. performed rhizobial growth assays. M.T. and D.M. analyzed the data and drafted the manuscript. M.Y., R.K.S and S.K. generated tissue and extracted RNA for chickpea transcriptome. M.T. processed the raw sequencing reads and quantified the gene expression levels in RNA-seq analysis. M.T., D.M., and J.M.A. provided important materials and guidance.

## Extended data

**Extended Data Fig. 1. Heat stress slows rhizobial growth and reduces *nod* gene induction. a,** Growth curves of *S. meliloti* 1021 at 28, 30, and 32°C monitored by OD600 for 96 h. Lines show the mean of 72 wells and shaded ribbons the standard deviation. **b-e,** Growth parameters extracted from Gompertz fits of each well: maximum growth rate (μ_max_; **b**), inverse time to mid-exponential phase (1/T_mid_; **c**), area under the curve (AUC; **d**), and carrying capacity (K; **e**). Different letters indicate statistically significant differences (Kruskal-Wallis test followed by Dunn’s test in **b**, **c**, and **e**; Welch’s ANOVA followed by the Games-Howell test in **d**; p-value < 0.05). **f,** Drop plate assay of *M. ciceri* CC1192 on YMB agar at 28, 30, and 32°C (R1-R3, three replicate spots). **g,** β-galactosidase activity (Miller units) of *S. meliloti* GMI5874 carrying a *nodA*pro::*lacZ* fusion after mock or luteolin treatment at 28, 30, and 32°C. Bars show the mean, error bars the standard deviation, and dots individual replicates (n = 8). Different letters indicate statistically significant differences (one-way ANOVA on log-transformed values followed by Dunnett’s T3 test; p-value < 0.05). **h,** Nodule number and **i,** nitrogenase activity measured by the ARA at 14 dpi in heat-stressed Medicago plants inoculated with *S. meliloti* 2011 (wild type, n = 14) or its *nodD*-overexpressing derivative 6390 (n = 23). Statistical significance was determined using two-sided Welch’s t-tests. Box plots show the median (center line), 25th and 75th percentiles (box), and whiskers extending to 1.5× the interquartile range, with individual data points shown as dots.

**Extended Data Fig. 2. Constitutive expression of the *NIN-LBD16-NFYA1-NFYB16* regulatory module rescues nodule organogenesis under heat stress. a,** Quantification of nodule number in plants constitutively expressing the *NIN-LBD16-NFYA1-NFYB16* module (n = 11) compared with the empty vector (n = 14) under control conditions, and **b,** heat stress (n = 12 (*NIN-LBD16-NFYA1-NFYB16*), n = 12 (WT-VC)). **c,** Nitrogenase activity measured by the ARA at 14 days post inoculation (dpi) under control (n = 10 (*NIN-LBD16-NFYA1-NFYB16*), n = 14 (WT-VC)) and **d,** heat stress (n = 12 (*NIN-LBD16-NFYA1-NFYB16*), n = 12 (WT-VC)). Statistical significance was determined using two-sided Welch’s t-tests. Box plots show the median (center line), 25th and 75th percentiles (box), and whiskers extending to 1.5× the interquartile range, with individual data points shown as dots.

**Extended Data Fig. 3. *M. truncatula* CEP peptides regulate root nodule symbiosis under heat stress. a,** Heatmap showing the expression patterns of 16 *M. truncatula CEP* genes under control and heat stress conditions at 0.5, 1, 3, 6, 24, and 48 hpi. **b,** Representative images of roots showing nodules on plants treated with CEP peptides under control or heat stress conditions. **c,** Shoot weight (mg) of Mock (n = 36), CEP1p (n = 72), CEP2p (n = 56), CEP5p (n = 50), CEP7p (n = 60), CEP8p (n = 62) and CEP12p (n = 56) treated plants under heat stress. **d,** Root weight (mg) of Mock (n = 36), CEP1p (n = 72), CEP2p (n = 56), CEP5p (n = 50), CEP7p (n = 60), CEP8p (n = 62) and CEP12p (n = 56) treated plants under heat stress. **e,** Root length (cm) of Mock (n = 36), CEP1p (n = 72), CEP2p (n = 56), CEP5p (n = 50), CEP7p (n = 60), CEP8p (n = 62) and CEP12p (n = 56) treated plants under heat stress. Statistical significance was determined using one-way ANOVA followed by Fisher’s LSD test (p-value < 0.05). Different letters indicate statistically significant differences. Box plots show the median (center line), 25th and 75th percentiles (box), and whiskers extending to 1.5× the interquartile range, with individual data points shown as dots.

**Extended Data Fig. 4. Autoregulation of nodulation pathway regulates root nodule symbiosis under heat stress. a,** Representative images of plants treated with synthetic CLE5 peptide at 1 μM compared to mock under control conditions. **b,** Quantification of the number of nodules in plants treated with CLE5 peptide compared to mock under control conditions (Mock (n = 13), MtCLE5p (n = 11)). **c,** Quantification of the number of nodules in plants constitutively expressing CLE5 compared to vector under control conditions (WT-VC (n = 30), *CLE5-*OE (n = 19)) and **d,** heat stress (WT-VC (n = 30), *CLE5-*OE (n = 32)). **e,** Quantification of the number of nodules in miR2111 overexpression plants compared to vector under control conditions (WT-VC (n = 15), *miR2111-*OE (n = 17)). Scale bar, 1 cm (**a**). Statistical significance was determined using two-sided Welch’s t-tests. Box plots show the median (center line), 25th and 75th percentiles (box), and whiskers extending to 1.5× the interquartile range, with individual data points shown as dots.

## Supplementary information

Supplementary Fig. 1. Gene ontology enrichment analysis of DEGs in *M. truncatula*.

Supplementary Fig. 2. Expression profiles of *CLE* genes in *M. truncatula*.

Supplementary Fig. 3. *M. truncatula* hypernodulation mutants show improved growth under heat stress.

Supplementary Fig. 4. Transcriptomic analysis of the effects of heat stress on root nodule symbiosis in *Cicer arietinum*.

Supplementary Fig. 5. Gene ontology enrichment analysis of DEGs in *Cicer arietinum*.

Supplementary Fig. 6. Expression profiles of *CEP* and *CLE* genes in *Cicer arietinum*.

Supplementary Table 1. Differentially expressed genes in *M. truncatula* under heat stress at 0.5, 1, 3, 6, 24, and 48 hpi (Excel file).

Supplementary Table 2. Differentially expressed genes in *C. arietinum* under heat stress at 0.5, 1, 3, 6, 24, and 48 hpi (Excel file).

Supplementary Table 3. CEP and CLE genes identified in the chickpea genome (Excel file).

Supplementary Table 4. Primers used in this study (Excel file).

## References

1. Martin, F. M., Uroz, S. & Barker, D. G. Ancestral alliances: Plant mutualistic symbioses with fungi and bacteria. Science 356, eaad4501 (2017).

2. Kundu, A., Moraes, T. A., Price, R. J., Harrison, R. J. & Oldroyd, G. E. D. Getting to the route: The evolution of nitrogen-fixing nodules. Annu. Rev. Cell Dev. Biol. 10.1146/annurev-cellbio-101123-093247 (2025) doi:10.1146/annurev-cellbio-101123-093247.

3. Radutoiu, S. et al. Plant recognition of symbiotic bacteria requires two LysM receptor-like kinases. Nature 425, 585–592 (2003).

4. Limpens, E. et al. LysM domain receptor kinases regulating rhizobial Nod factor-induced infection. Science 302, 630–633 (2003).

5. Stracke, S. et al. A plant receptor-like kinase required for both bacterial and fungal symbiosis. Nature 417, 959–962 (2002).

6. Endre, G. et al. A receptor kinase gene regulating symbiotic nodule development. Nature 417, 962–966 (2002).

7. Singh, S., Katzer, K., Lambert, J., Cerri, M. & Parniske, M. CYCLOPS, a DNA-binding transcriptional activator, orchestrates symbiotic root nodule development. Cell Host Microbe 15, 139–152 (2014).

8. Yano, K. et al. CYCLOPS, a mediator of symbiotic intracellular accommodation. Proc. Natl. Acad. Sci. U. S. A. 105, 20540–20545 (2008).

9. Tirichine, L. et al. Deregulation of a Ca2+/calmodulin-dependent kinase leads to spontaneous nodule development. Nature 441, 1153–1156 (2006).

10. Lévy, J. et al. A putative Ca2+ and calmodulin-dependent protein kinase required for bacterial and fungal symbioses. Science 303, 1361–1364 (2004).

11. Oldroyd, G. E. D. & Downie, J. A. Coordinating nodule morphogenesis with rhizobial infection in legumes. Annu. Rev. Plant Biol. 59, 519–546 (2008).

12. Liu, C.-W. et al. NIN Acts as a Network Hub Controlling a Growth Module Required for Rhizobial Infection. Plant Physiol. 179, 1704–1722 (2019).

13. Cathebras, C. et al. A novel cis-element enabled bacterial uptake by plant cells. Nat. Plants 12, 140–151 (2026).

14. Liu, J. et al. Ancestral functionality and symbiotic refinement of NIN in root nodule symbiosis. Nat. Commun. 1–11 (2026).

15. Tiwari, M. et al. Evolutionary and functional analysis of two-component system in chickpea reveals CaRR13, a TypeB RR, as positive regulator of symbiosis. Plant Biotechnol. J. 19, 2415–2427 (2021).

16. Gonzalez-Rizzo, S., Crespi, M. & Frugier, F. The Medicago truncatula CRE1 cytokinin receptor regulates lateral root development and early symbiotic interaction with Sinorhizobium meliloti. Plant Cell 18, 2680–2693 (2006).

17. Soyano, T., Shimoda, Y., Kawaguchi, M. & Hayashi, M. A shared gene drives lateral root development and root nodule symbiosis pathways in Lotus. Science 366, 1021–1023 (2019).

18. Schiessl, K. et al. NODULE INCEPTION Recruits the Lateral Root Developmental Program for Symbiotic Nodule Organogenesis in Medicago truncatula. Curr. Biol. 29, 3657–3668.e5 (2019).

19. Laffont, C. & Frugier, F. Rhizobium symbiotic efficiency meets CEP signaling peptides. New Phytol. 241, 24–27 (2024).

20. Gresshoff, P. M. et al. Functional genomics dissection of the nodulation autoregulation pathway (AON) in soybean (Glycine max). J. Integr. Plant Biol. 67, 762–772 (2025).

21. Schnabel, E., Journet, E.-P., de Carvalho-Niebel, F., Duc, G. & Frugoli, J. The Medicago truncatula SUNN gene encodes a CLV1-like leucine-rich repeat receptor kinase that regulates nodule number and root length. Plant Mol. Biol. 58, 809–822 (2005).

22. Schnabel, E. et al. The lss supernodulation mutant of Medicago truncatula reduces expression of the SUNN gene. Plant Physiol. 154, 1390–1402 (2010).

23. Schnabel, E. L. et al. The ROOT DETERMINED NODULATION1 gene regulates nodule number in roots of Medicago truncatula and defines a highly conserved, uncharacterized plant gene family. Plant Physiol. 157, 328–340 (2011).

24. Schnabel, E. et al. The Defective in Autoregulation (DAR) gene of Medicago truncatula encodes a protein involved in regulating nodulation and arbuscular mycorrhiza. BMC Plant Biol. 24, 766 (2024).

25. Smith, S. et al. The CEP5 peptide promotes abiotic stress tolerance, as revealed by quantitative proteomics, and attenuates the AUX/IAA equilibrium in Arabidopsis. Mol. Cell. Proteomics 19, 1248–1262 (2020).

26. Wu, S. et al. Soybean CEP6 signaling peptides positively regulate nodulation. Agronomy (Basel) 14, 988 (2024).

27. Räsänen, L. A., Elväng, A. M., Jansson, J. & Lindström, K. Effect of heat stress on cell activity and cell morphology of the tropical rhizobium, Sinorhizobium arboris. FEMS Microbiol. Ecol. 34, 267–278 (2001).

28. Munévar, F. & Wollum, A. G., II. Growth of Rhizobium japonicum Strains at Temperatures Above 27°C. Appl. Environ. Microbiol. 42, 272–276 (1981).

29. Karanja, N. K. & Wood, M. SelectingRhizobium phaseoli strains for use with beans (Phaseolus vulgaris L.) in Kenya: Tolerance of high temperature and antibiotic resistance. Plant Soil 112, 15–22 (1988).

30. Räsänen, L. A. & Lindströmm, K. The effect of heat stress on the symbiotic interaction between Sinorhizobium sp. and Acacia senegal. FEMS Microbiol. Ecol. 28, 63–74 (1999).

31. Räsänen, L. A. & Lindström, K. The effect of heat stress on the symbiotic interaction between Sinorhizobium sp. and Acacia senegal. FEMS Microbiol. Ecol. 28, 63–74 (1999).

32. Serova, T. A., et al. Effects of elevated temperature on Pisum sativum nodule development: I-detailed characteristic of unusual apical senescence. Int. J. Mol. Sci. 24, 17144 (2023).

33. Kitaeva, A. B., Serova, T. A., Kusakin, P. G. & Tsyganov, V. E. Effects of elevated temperature on Pisum sativum nodule development: II-phytohormonal responses. Int. J. Mol. Sci. 24, 17062 (2023).

34. Soba, D., Arrese-Igor, C. & Aranjuelo, I. Additive effects of heatwave and water stresses on soybean seed yield is caused by impaired carbon assimilation at pod formation but not at flowering. Plant Sci. 321, 111320 (2022).

35. Hernandez-Armenta, R., Wien, H. C. & Eaglesham, A. R. J. Carbohydrate partitioning and nodule function in common bean after heat stress. Crop Sci. 29, 1292–1297 (1989).

36. Serova, T. A., et al. Effects of elevated temperature on Pisum sativum nodule development: I-detailed characteristic of unusual apical senescence. Int. J. Mol. Sci. 24, 17144 (2023).

37 Kitaeva, A. B., Serova, T. A., Kusakin, P. G. & Tsyganov, V. E. Effects of elevated temperature on Pisum sativum nodule development: II-phytohormonal responses. Int. J. Mol. Sci. 24, 17062 (2023).

38. Munns, D. N., Fogle, V. W. & Hallock, B. G. Alfalfa root nodule distribution and inhibition of nitrogen fixation by heat^1^. Agron. J. 69, 377–380 (1977).

39. Kalil, A. et al. NUP107-160 nuclear pore sub-complex members determine symbiotic ion channel localization in legumes. Plant Physiol. kiag415 (2026).

40. Ferguson, B. J. et al. Legume nodulation: The host controls the party. Plant Cell Environ. 42, 41–51 (2019).

41. Hungria, M. & Vargas, M. A. T. Environmental factors affecting N2 fixation in grain legumes in the tropics, with an emphasis on Brazil. Field Crops Res. 65, 151–164 (2000).

42. Soyano, T., Kouchi, H., Hirota, A. & Hayashi, M. Nodule inception directly targets NF-Y subunit genes to regulate essential processes of root nodule development in Lotus japonicus. PLoS Genet. 9, e1003352 (2013).

43. Gautrat, P., Laffont, C. & Frugier, F. Compact Root Architecture 2 Promotes Root Competence for Nodulation through the miR2111 Systemic Effector. Curr. Biol. 30, 1339–1345.e3 (2020).

44. Taleski, M. et al. CEP hormones at the nexus of nutrient acquisition and allocation, root development, and plant-microbe interactions. J. Exp. Bot. 75, 538–552 (2024).

45. Isidra-Arellano, M. C. et al. Inhibition of legume nodulation by Pi deficiency is dependent on the autoregulation of nodulation (AON) pathway. Plant J. 103, 1125–1139 (2020).

46. Maillet, F., Debellé, F. & Dénarié, J. Role of the nodD and syrM genes in the activation of the regulatory gene nodD3, and of the common and host-specific nod genes of Rhizobium meliloti. Mol. Microbiol. 4, 1975–1984 (1990).

47. Trimborn, L., Hoecker, U. & Ponnu, J. A simple quantitative assay for measuring β-galactosidase activity using X-gal in yeast-based interaction analyses. Curr. Protoc2, e421 (2022).

48. Gentile, I., Santo Domingo, M., Zebell, S. G., Fitzgerald, B. & Lippman, Z. B. Pan-angiosperm analysis of the CLE signaling peptide gene family unveils paths, patterns, and predictions of paralog diversification. Mol. Biol. Evol. 42, msaf294 (2025).

49. Pimentel, H., Bray, N. L., Puente, S., Melsted, P. & Pachter, L. Differential analysis of RNA-seq incorporating quantification uncertainty. Nat. Methods 14, 687–690 (2017).

50. Tian, F., Yang, D.-C., Meng, Y.-Q., Jin, J. & Gao, G. PlantRegMap: charting functional regulatory maps in plants. Nucleic Acids Res. 48, D1104–D1113 (2020).

51. Patron, N. J. et al. Standards for plant synthetic biology: a common syntax for exchange of DNA parts. New Phytol. 208, 13–19 (2015).

52. Catoira, R. et al. Four genes of Medicago truncatula controlling components of a nod factor transduction pathway. Plant Cell 12, 1647 (2000).

53. Zhao, L., Ji, C.-Y., Murray, J. D. & Liu, C.-W. A legume cellulase required for rhizobial infection and colonization in root nodule symbiosis. Nat. Commun. 16, 6663 (2025).

54. Starker, C. G., Parra-Colmenares, A. L., Smith, L., Mitra, R. M. & Long, S. R. Nitrogen fixation mutants of Medicago truncatula fail to support plant and bacterial symbiotic gene expression. Plant Physiol. 140, 671–680 (2006).

55. Schmitz, D. A., Wechsler, T., Mignot, I. & Kümmerli, R. Predicting bacterial interaction outcomes from monoculture growth and supernatant assays. ISME Commun. 4, ycae045 (2024).

