## Extended Fig 1-4 for "Rewiring systemic nodulation control to engineer heat-tolerant nitrogen fixation in *Medicago truncatula* and chickpea"

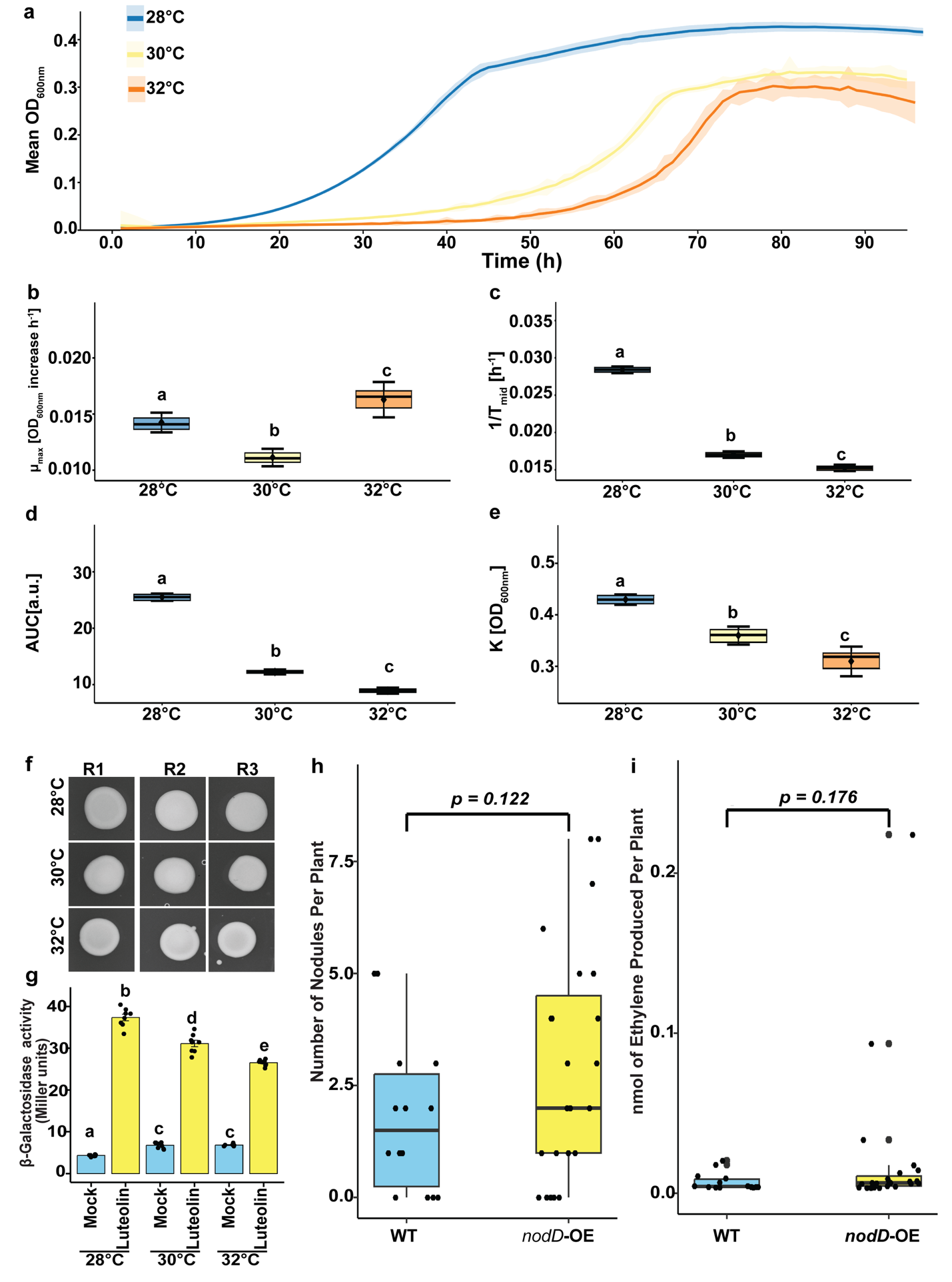

**Extended Data Fig. 1 | Heat stress slows rhizobial growth and reduces *nod* gene induction. a,** Growth curves of *S. meliloti* 1021 at 28, 30, and 32°C monitored by OD600 for 96 h. Lines show the mean of 72 wells and shaded ribbons the standard deviation. **b-e,** Growth parameters extracted from Gompertz fits of each well: maximum growth rate (μ_max_; **b**), inverse time to mid-exponential phase (1/T_mid_; **c**), area under the curve (AUC; **d**), and carrying capacity (K; **e**). Different letters indicate statistically significant differences (Kruskal-Wallis test followed by Dunn's test in **b**, **c**, and **e**; Welch's ANOVA followed by the Games-Howell test in **d**; p-value < 0.05). **f,** Drop plate assay of *M. ciceri* CC1192 on YMB agar at 28, 30, and 32°C (R1-R3, three replicate spots). **g,** β-galactosidase activity (Miller units) of *S. meliloti* GMI5874 carrying a *nodA*pro::*lacZ* fusion after mock or luteolin treatment at 28, 30, and 32°C. Bars show the mean, error bars the standard deviation, and dots individual replicates (n = 8). Different letters indicate statistically significant differences (one-way ANOVA on log-transformed values followed by Dunnett's T3 test; p-value < 0.05). **h,** Nodule number and **i,** nitrogenase activity measured by the ARA at 14 dpi in heat-stressed Medicago plants inoculated with *S. meliloti* 2011 (wild type, n = 14) or its *nodD*-overexpressing derivative 6390 (n = 23). Statistical significance was determined using two-sided Welch's t-tests. Box plots show the median (center line), 25th and 75th percentiles (box), and whiskers extending to 1.5× the interquartile range, with individual data points shown as dots.

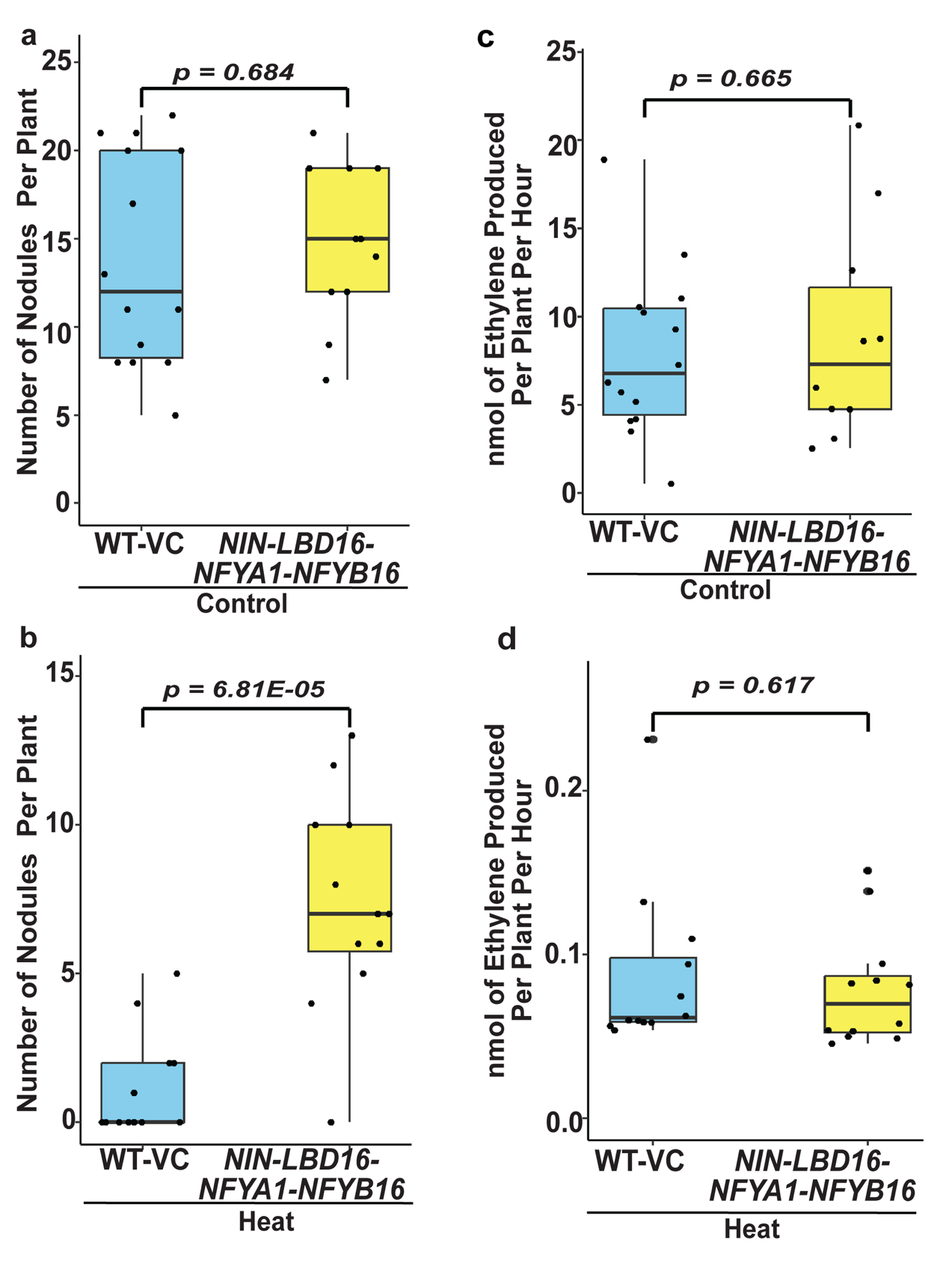

**Extended Data Fig. 2 | Constitutive expression of the *NIN-LBD16-NFYA1-NFYB16* regulatory module rescues nodule organogenesis under heat stress. a,** Quantification of nodule number in plants constitutively expressing the *NIN-LBD16-NFYA1-NFYB16* module (n = 11) compared with the empty vector (n = 14) under control conditions, and **b,** heat stress (n = 12 (*NIN-LBD16-NFYA1-NFYB16*), n = 12 (WT-VC)). **c,** Nitrogenase activity measured by the ARA at 14 days post inoculation (dpi) under control (n = 10 (*NIN-LBD16-NFYA1-NFYB16*), n = 14 (WT-VC)) and **d,** heat stress (n = 12 (*NIN-LBD16-NFYA1-NFYB16*), n = 12 (WT-VC)). Statistical significance was determined using two-sided Welch's t-tests. Box plots show the median (center line), 25th and 75th percentiles (box), and whiskers extending to 1.5× the interquartile range, with individual data points shown as dots.

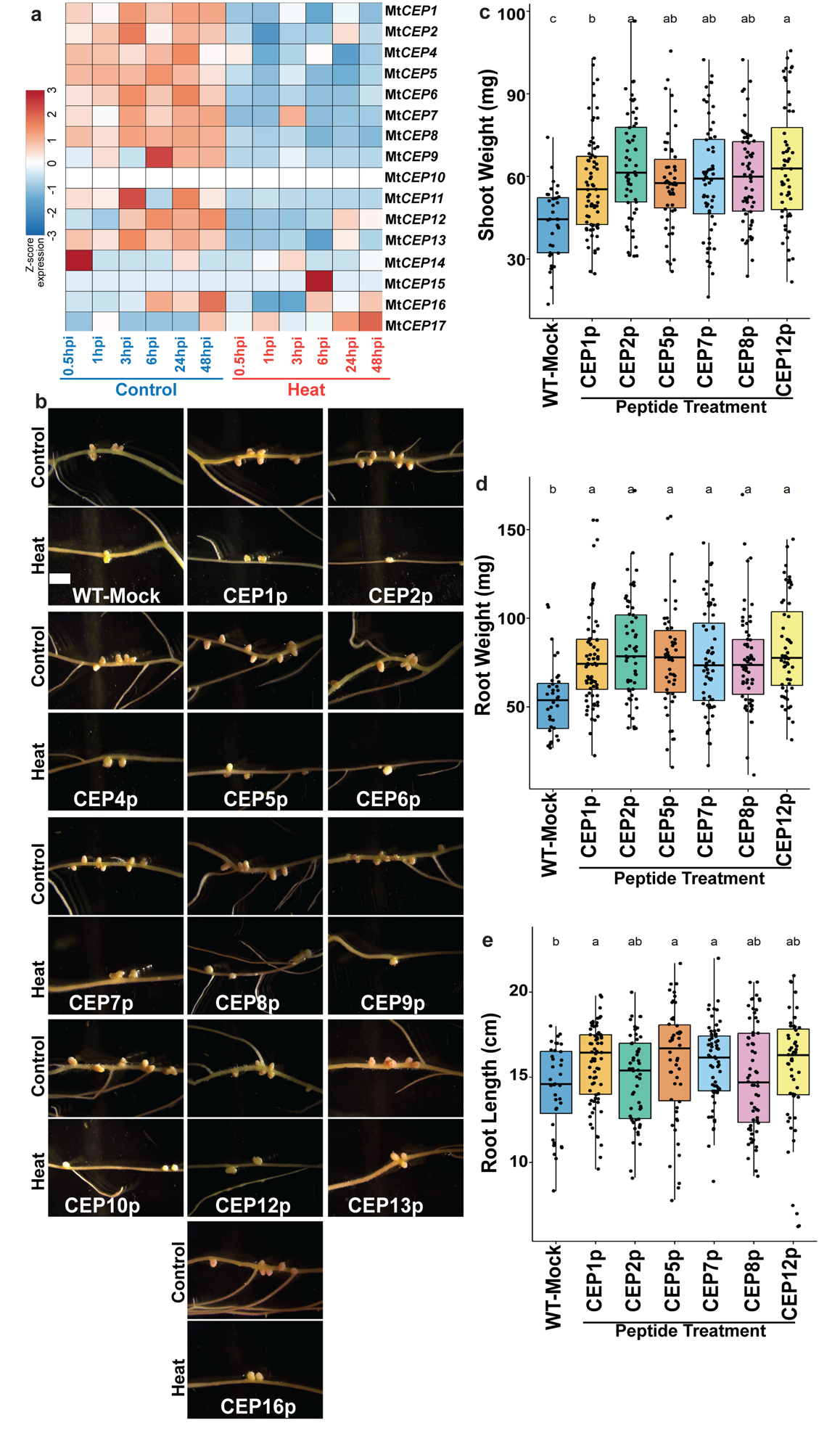

**Extended Data Fig. 3 | *M. truncatula* CEP peptides regulate root nodule symbiosis under heat stress. a,** Heatmap showing the expression patterns of 16 *M. truncatula* *CEP* genes under control and heat stress conditions at 0.5, 1, 3, 6, 24, and 48 hpi. **b,** Representative images of roots showing nodules on plants treated with CEP peptides under control or heat stress conditions. **c,** Shoot weight (mg) of Mock (n = 36), CEP1p (n = 72), CEP2p (n = 56), CEP5p (n = 50), CEP7p (n = 60), CEP8p (n = 62) and CEP12p (n = 56) treated plants under heat stress. **d,** Root weight (mg) of Mock (n = 36), CEP1p (n = 72), CEP2p (n = 56), CEP5p (n = 50), CEP7p (n = 60), CEP8p (n = 62) and CEP12p (n = 56) treated plants under heat stress. **e,** Root length (cm) of Mock (n = 36), CEP1p (n = 72), CEP2p (n = 56), CEP5p (n = 50), CEP7p (n = 60), CEP8p (n = 62) and CEP12p (n = 56) treated plants under heat stress. Statistical significance was determined using one-way ANOVA followed by Fisher's LSD test (p-value < 0.05). Different letters indicate statistically significant differences. Box plots show the median (center line), 25th and 75th percentiles (box), and whiskers extending to 1.5× the interquartile range, with individual data points shown as dots.

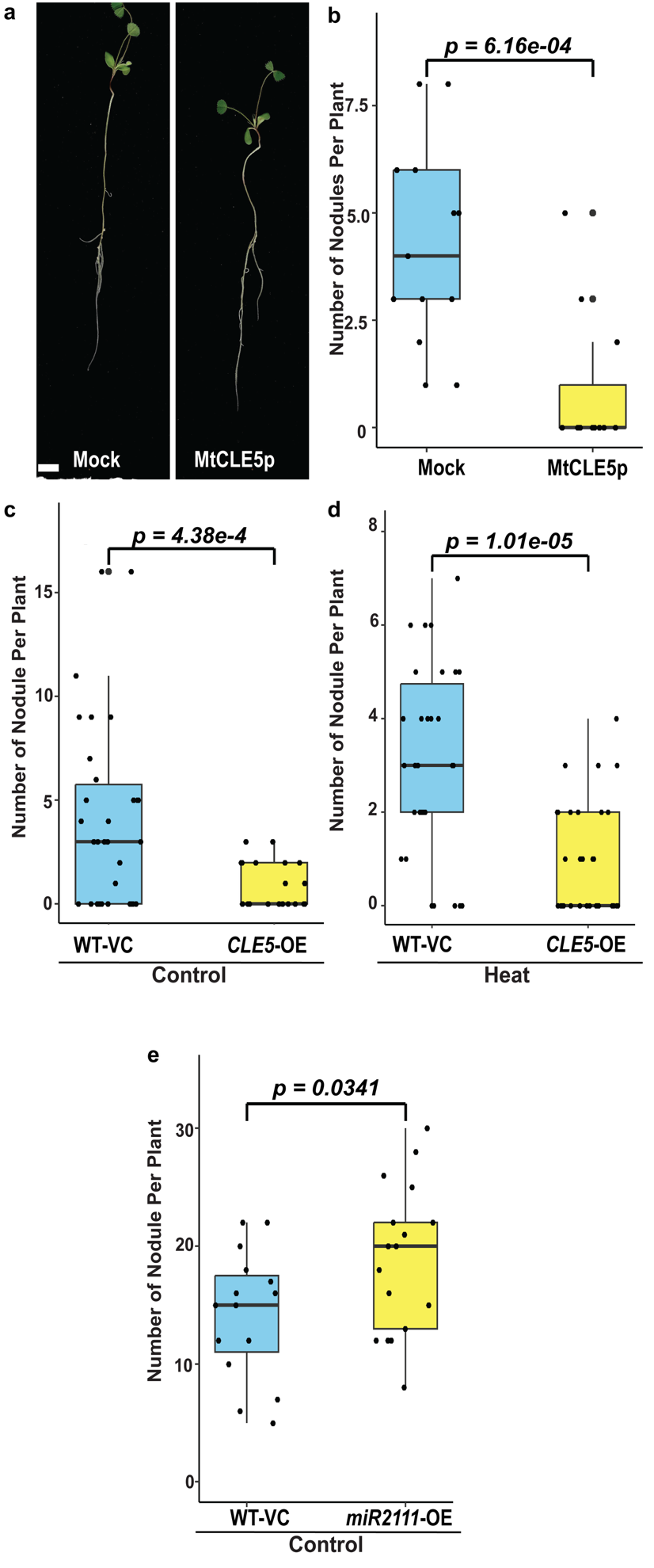

**Extended Data Fig. 4 | Autoregulation of nodulation pathway regulates root nodule symbiosis under heat stress. a,** Representative images of plants treated with synthetic CLE5 peptide at 1 μM compared to mock under control conditions. **b,** Quantification of the number of nodules in plants treated with CLE5 peptide compared to mock under control conditions (Mock (n = 13), MtCLE5p (n = 11)). **c,** Quantification of the number of nodules in plants constitutively expressing CLE5 compared to vector under control conditions (WT-VC (n = 30), *CLE5-*OE (n = 19)) and **d,** heat stress (WT-VC (n = 30), *CLE5-*OE (n = 32)). **e,** Quantification of the number of nodules in miR2111 overexpression plants compared to vector under control conditions (WT-VC (n = 15), *miR2111-*OE (n = 17)). Scale bar, 1 cm (**a**). Statistical significance was determined using two-sided Welch's t-tests. Box plots show the median (center line), 25th and 75th percentiles (box), and whiskers extending to 1.5× the interquartile range, with individual data points shown as dots.
