## Supplementary Fig 1-6 for "Rewiring systemic nodulation control to engineer heat-tolerant nitrogen fixation in *Medicago truncatula* and chickpea"

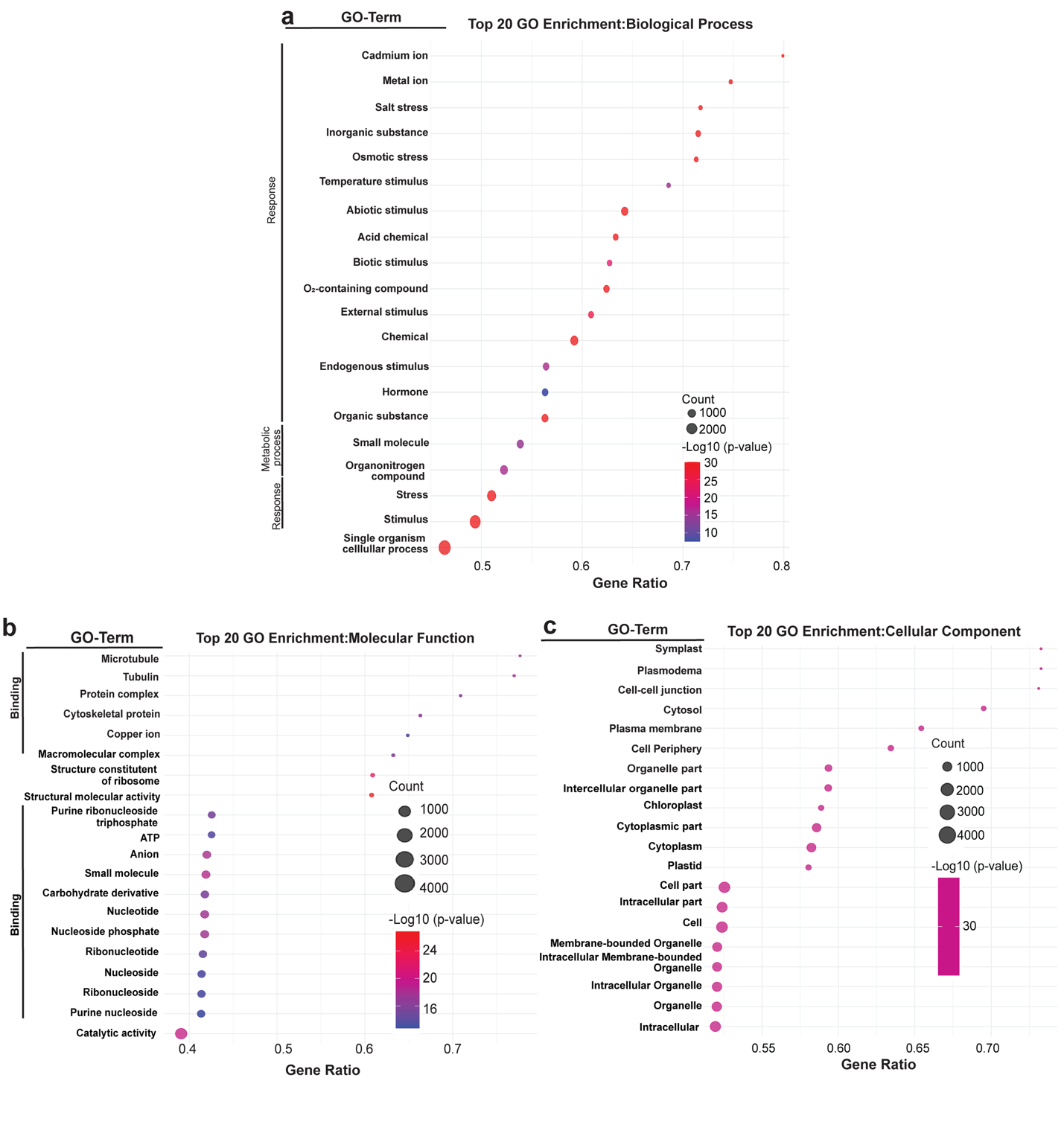

**Supplementary Fig. 1 | Gene ontology (GO) enrichment analysis of DEGs in *M. truncatula*. a,** Gene Ontology (GO) enrichment analysis of *M. truncatula* DEGs under heat stress compared with the control. Bubble plots show the top 20 enriched biological processes. **b,** molecular function. **c,** cellular component. Bubble size indicates the number of genes associated with each GO term, and bubble color represents the enrichment significance (-log10 p-value). Gene ratio on the x-axis represents the DEGs annotated in a GO term with respect to total DEGs. GO terms are classified broadly into “Response” and “Metabolic process” on the y-axis in **a**, and "Binding" in **b**. A Wald test was used to identify DEGs between heat stress and control conditions at 0.5, 1, 3, 6, 24, and 48 hours post-inoculation (hpi), with statistical significance set at p-value ≤ 0.05 and false-discovery rate (q-value) ≤ 0.01. GO enrichment analysis was performed using the Plant Transcriptional Regulatory Map database, which applies topGO and Fisher's exact tests to determine significantly enriched GO terms at a threshold of p-value ≤ 0.01.

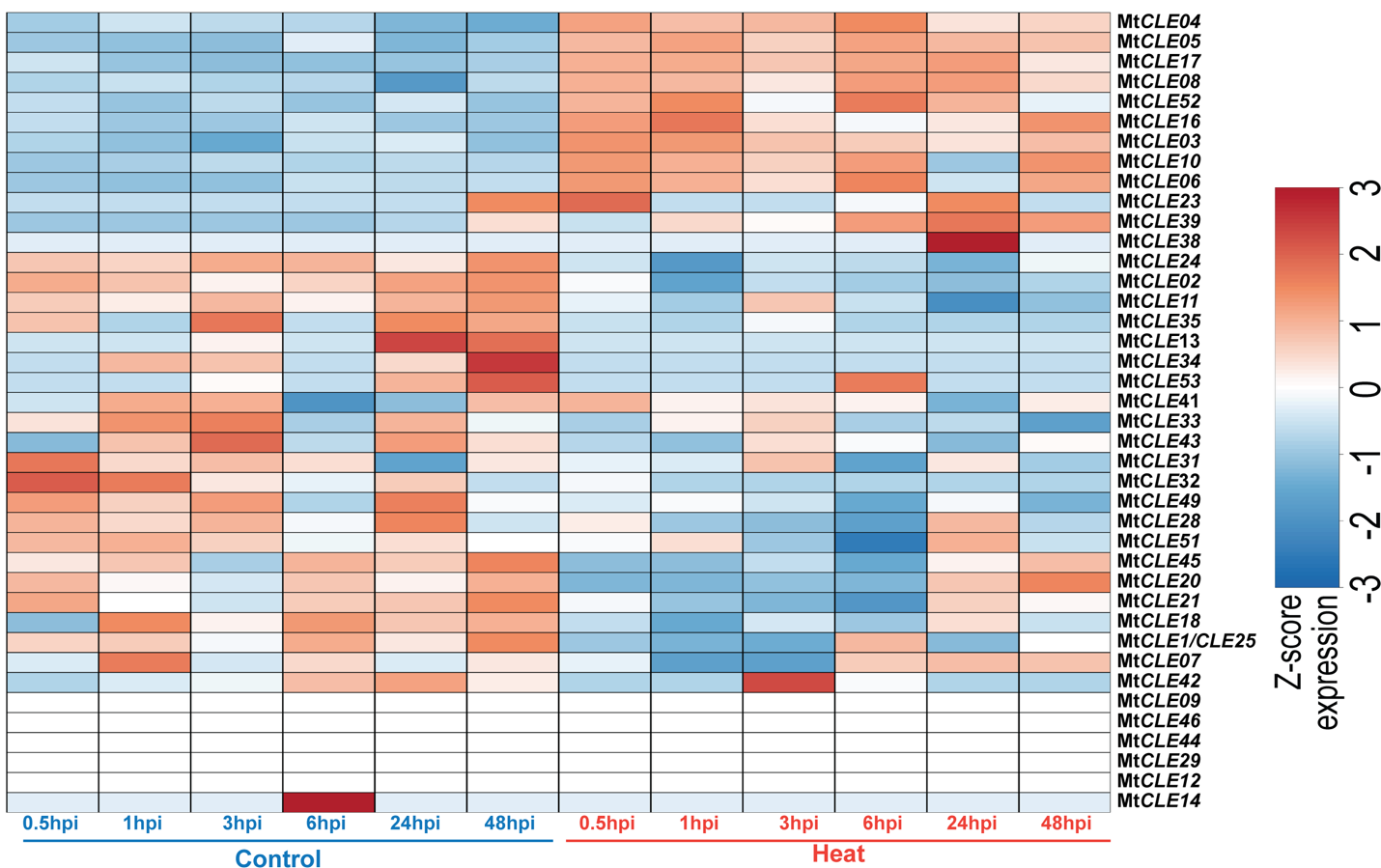

**Supplementary Fig. 2 | Expression profiles of *CLE* genes in *M. truncatula*.** Heatmap showing CLE gene expression profiles under control and heat stress conditions at 0.5, 1, 3, 6, 24, and 48 hpi. Data were obtained through RNA-seq of *Medicago* roots inoculated with *S. meliloti* 1021, in three independent biological replicates under control and heat stress conditions at different time points. The color scale represents row-scaled Z-scores of log2(TPM + 1)- transformed gene expression, clipped to a range of −3 to 3.

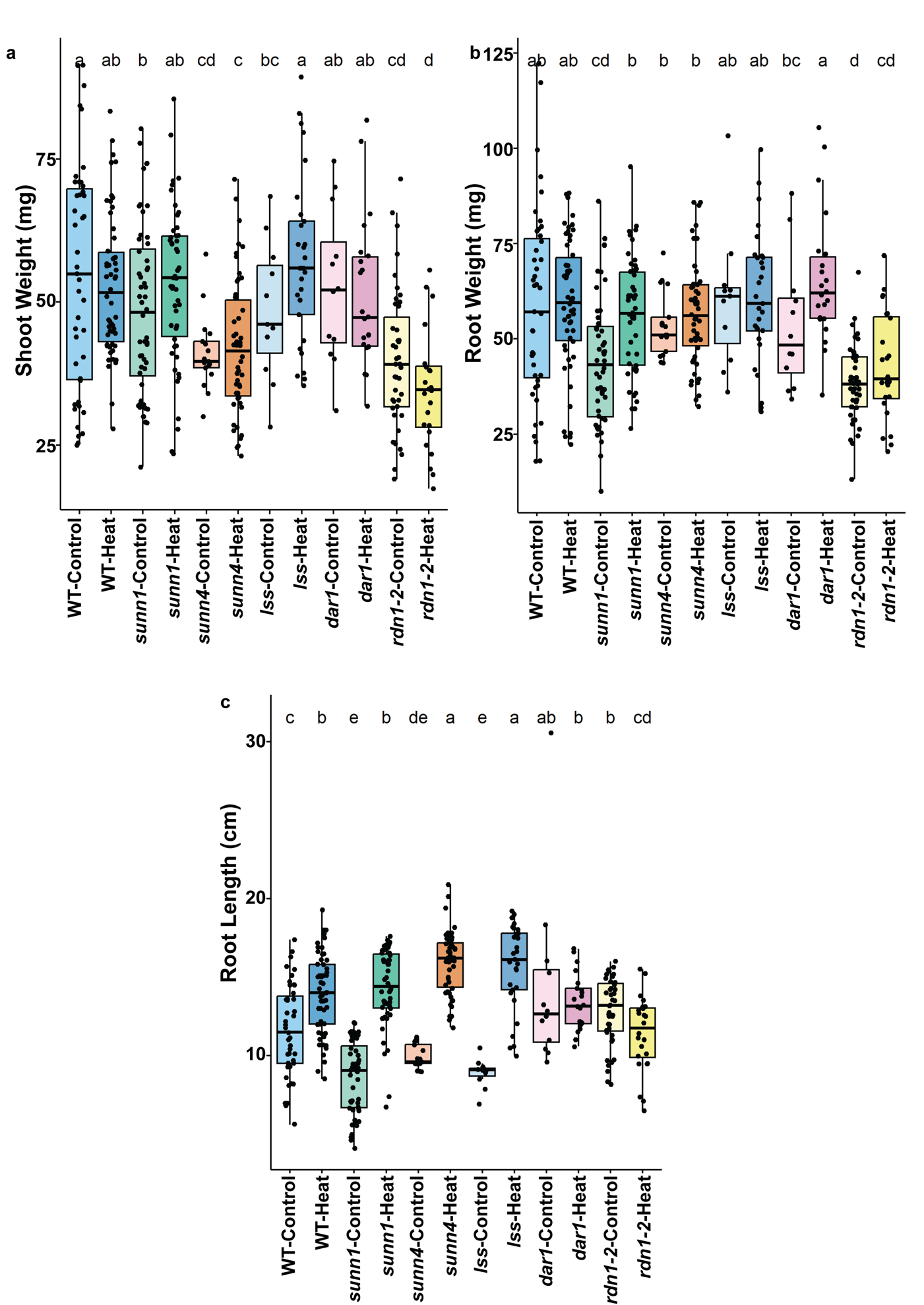

**Supplementary Fig. 3 | *Medicago truncatula* hypernodulation mutants show improved growth under heat stress. a,** Shoot weight (mg) of wild-type (WT), *sunn-1, sunn-4, lss, dar-1,* and *rdn1-2* under control and heat stress (WT-C (n = 43), WT-H (n = 54), *sunn-1*-C (n = 52), *sunn-1*-H (n = 50), *sunn-4*-C (n = 17), *sunn-4*-H (n = 50), *lss*-C (n = 11), *lss*-H (n = 29), *dar-1*-C (n = 12), *dar-1*-H (n = 22), *rdn1-2*-C (n = 43), *rdn1-2*-H (n = 24)). **b,** Root weight (mg) of WT, *sunn-1, sunn-4, lss, dar-1,* and *rdn1-2* under control and heat stress (WT-C (n = 43), WT-H (n = 54), *sunn-1*-C (n = 52), *sunn-1*-H (n = 50), *sunn-4*-C (n = 17), *sunn-4*-H (n = 50), *lss*-C (n = 11), *lss*-H (n = 29), *dar-1*-C (n = 12), *dar-1*-H (n = 22), *rdn1-2*-C (n = 43), *rdn1-2*-H (n = 24)). **c,** Root length (cm) of wild-type, *sunn-1, sunn-4, lss, dar-1,* and *rdn1-2* under control and heat stress (WT-C (n = 43), WT-H (n = 54), *sunn-1*-C (n = 52), *sunn-1*-H (n = 50), *sunn-4*-C (n = 17), *sunn-4*-H (n = 50), *lss*-C (n = 11), *lss*-H (n = 29), *dar-1*-C (n = 12), *dar-1*-H (n = 22), *rdn1-2*-C (n = 43), *rdn1-2*-H (n = 24)). Statistical significance was determined using one-way ANOVA followed by Fisher's LSD test (p-value < 0.05). Different letters indicate statistically significant differences. Box plots show the median (center line), 25th and 75th percentiles (box), and whiskers extending to 1.5× the interquartile range, with individual data points shown as dots.

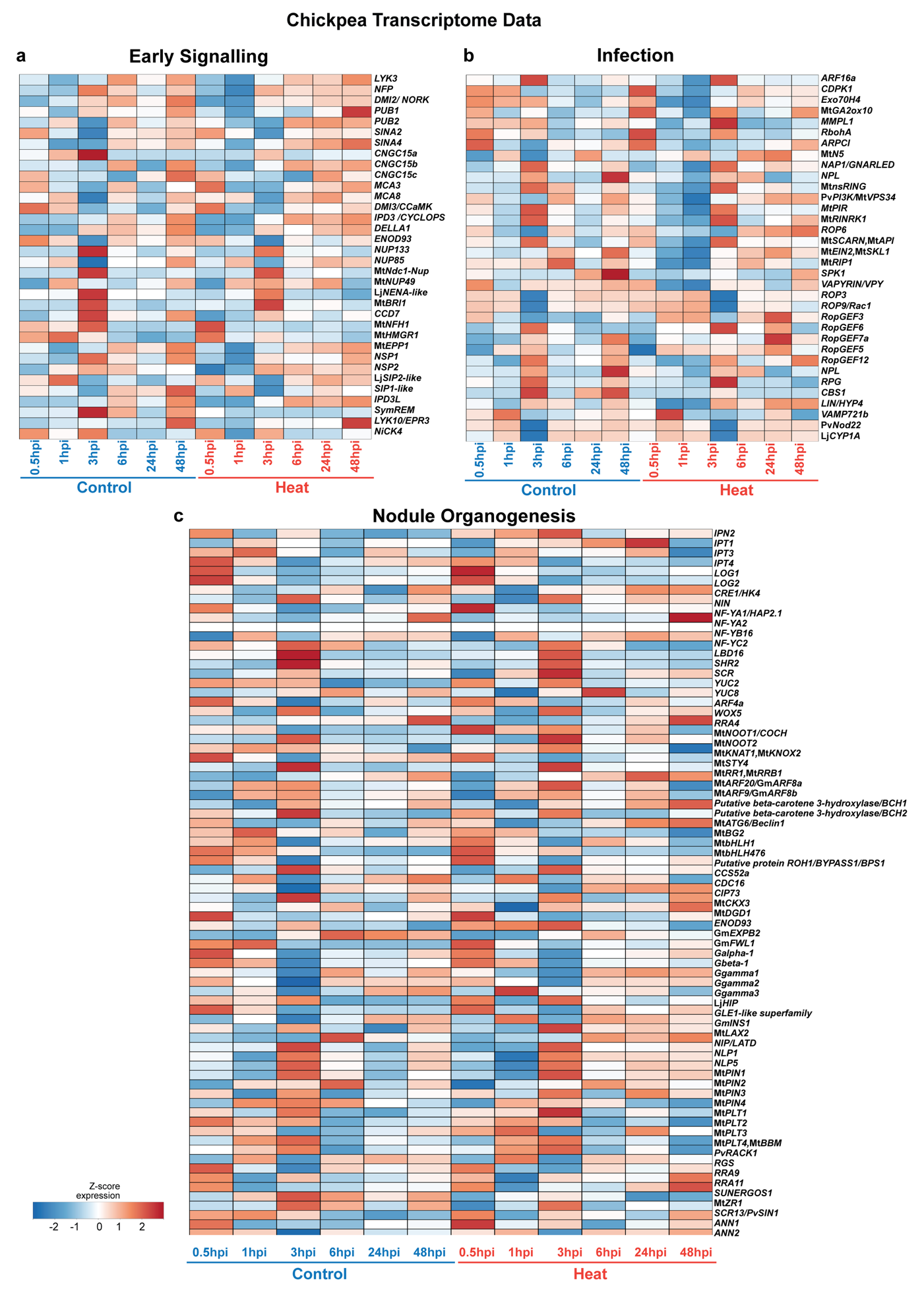

**Supplementary Fig. 4 | Transcriptomic analysis of the effects of heat stress on root nodule symbiosis in *Cicer arietinum* (chickpea). a,** Heatmaps showing the expression profiles of genes differentially expressed under control and heat stress conditions at 0.5, 1, 3, 6, 24, and 48 hours post-inoculation (hpi). Genes are grouped according to their roles in early symbiotic signaling, **b,** infection and **c,** nodule organogenesis. Data were obtained by RNA-seq of chickpea roots inoculated with *Mesorhizobium ciceri* CC1192, in three independent biological replicates under control and heat-stress conditions at different time points. Color scale represents row-scaled Z-score of log2 (TPM + 1) transformed gene expression clipped to a range of −2 to 2 shared across panels a- c.

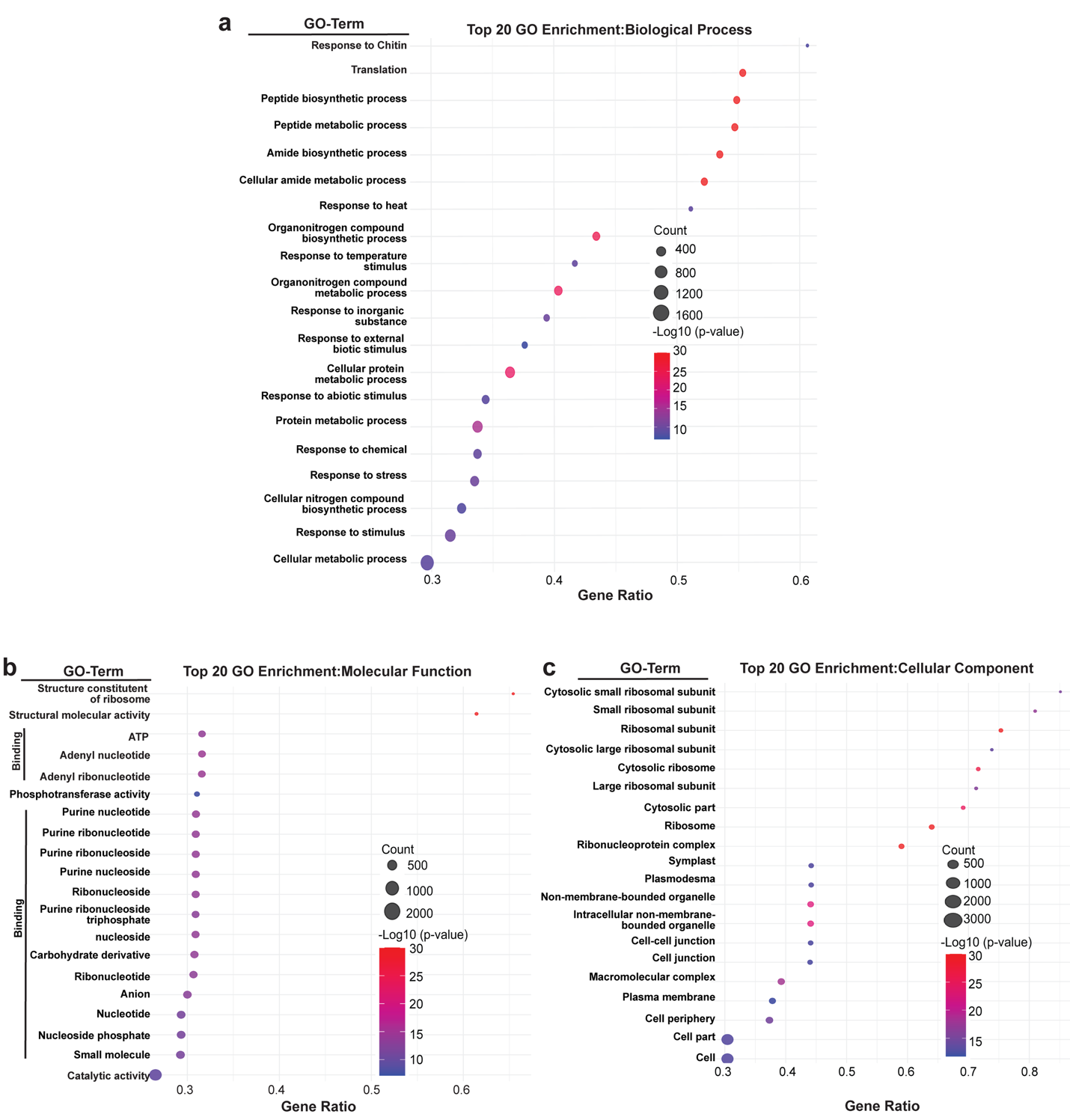

**Supplementary Fig. 5 | Gene ontology (GO) enrichment analysis of DEGs in *Cicer arietinum* (chickpea). a,** Gene Ontology (GO) enrichment analysis of chickpea DEGs under heat stress compared with the control. Bubble plots show the top 20 enriched biological processes. **b,** molecular function. **c,** cellular component. Bubble size indicates the number of genes associated with each GO term, and bubble color represents the enrichment significance (-log10 p-value). Gene ratio on the x-axis represents the DEGs annotated in a GO term with respect to total DEGs. GO terms are broadly classified as "Binding" in **b**. A Wald test was used to identify DEGs between heat stress and control conditions at 0.5, 1, 3, 6, 24, and 48 hours post-inoculation (hpi), with statistical significance set at p-value ≤ 0.05 and false-discovery rate (q-value) ≤ 0.01. GO enrichment analysis was performed using the Plant Transcriptional Regulatory Map database, which applies topGO and Fisher's exact tests to determine significantly enriched GO terms at a threshold of p-value ≤ 0.01.

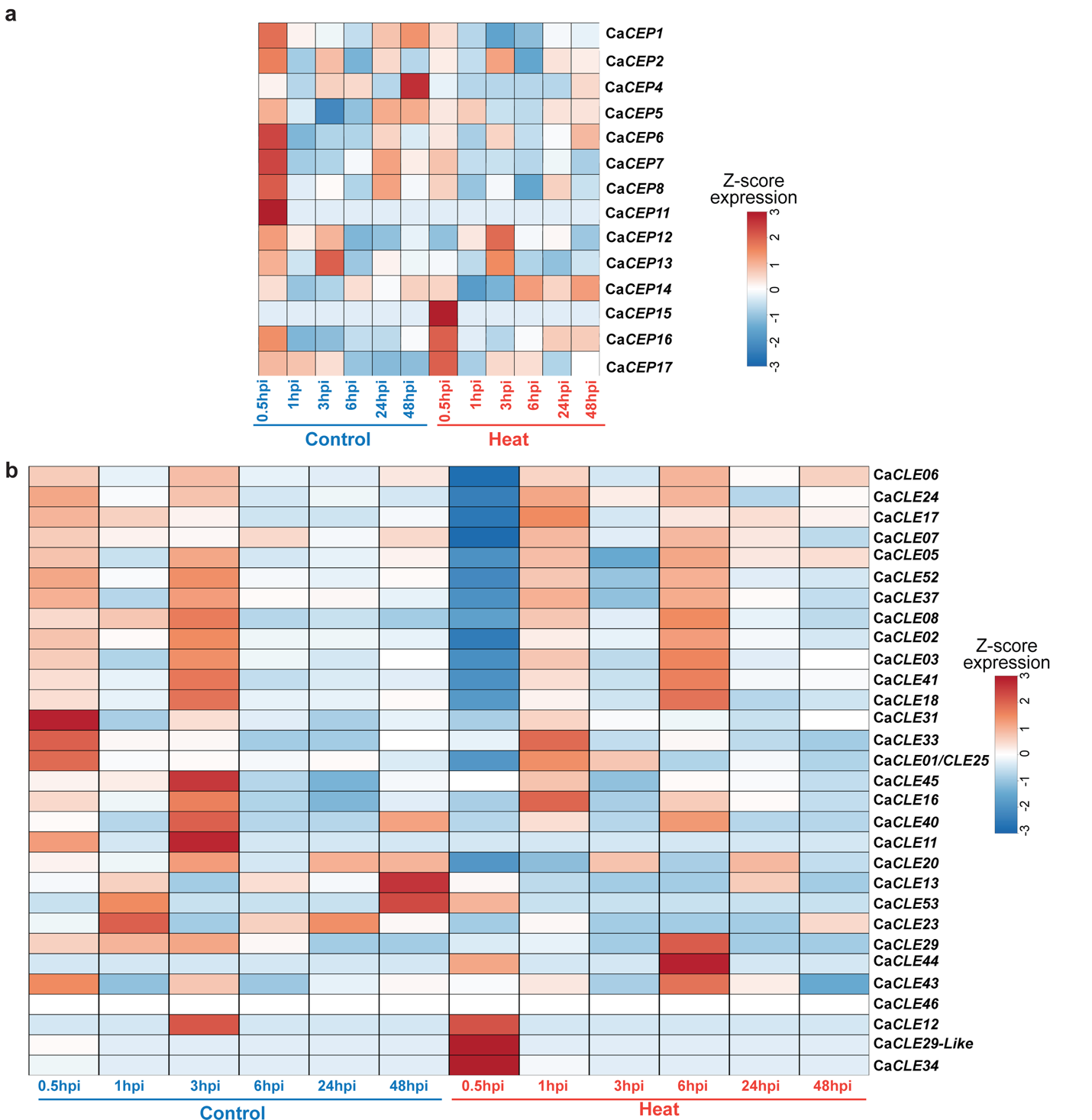

**Supplementary Fig. 6 | Expression profiles of *CEP* and *CLE* genes in *Cicer arietinum* (chickpea). a,** Heatmaps showing the expression profiles under control and heat stress conditions at 0.5, 1, 3, 6, 24, and 48 hpi of *CEP* genes, and **b,** *CLE* genes. Data were obtained through RNA-seq of chickpea roots inoculated with *Mesorhizobium ciceri* CC1192, in three independent biological replicates, under control and heat stress conditions at different time points. The color scale shows row-scaled Z-scores of log2(TPM + 1)- transformed gene expression, clipped to −3 to 3 and shared across panels a-b.
